# SILCS-Guided Feature Encoding Expands Ligand Recognition at an HBV Core Protein Interface

**DOI:** 10.64898/2026.08.16.745131

**Authors:** Zixing Fan, Ruoqing Jia, Diane L. Lynch, Anna Pavlova, Andrew C. McShan, James C. Gumbart

## Abstract

Hepatitis B virus (HBV) infection depends on coordinated capsid assembly and virion production. A hydrophobic pocket at the intradimer interface of the HBV core protein has been linked to secretion phenotypes and shown to bind small molecules, but its interaction landscape remains poorly defined. Here, we combine atomistic molecular dynamics (MD) simulations, fragment mapping, pharmacophore modeling, and biophysical binding assays to characterize this pocket and identify ligands with binding modes that extend beyond the known pocket. This workflow narrowed an initial library of approximately 4.7 million compounds to eight candidates for experimental evaluation by saturation transfer difference (STD) NMR and surface plasmon resonance (SPR). Of these eight, compound B3 showed detectable STD NMR signals, concentration-dependent SPR binding, and an MD-supported binding mode that retained hydrophobic-pocket anchoring while also extending toward the spike-proximal loop. Together, these results illustrate how dynamic fragment mapping can identify ligand candidates that engage broader interaction landscapes at protein-interface pockets.

## 1 Introduction

Chronic hepatitis B virus (HBV) infection remains a major global health burden, affecting approximately 240 million people worldwide, with mortality driven primarily by cirrhosis and hepatocellular carcinoma.^1,2^ Prophylactic vaccination has substantially reduced incident infections; however, incomplete vaccine uptake permits ongoing transmission, while the large reservoir of chronic infections continues to sustain disease burden.^1,3^ Current therapeutic strategies, primarily long-term nucleos(t)ide analogues, effectively suppress viral replication but rarely achieve a durable functional cure. This limitation arises in part from the persistence of covalently closed circular DNA and viral–host interactions that are not eliminated by polymerase inhibition.^4–7^ In addition, the potential emergence of drug-resistant viral variants during prolonged therapy can compromise long-term antiviral control.^8^ These challenges motivate the development of therapeutic approaches targeting additional stages of the HBV life cycle, including capsid assembly, nucleocapsid maturation, and virion envelopment and secretion.^9–12^

The HBV capsid is a metastable protein shell assembled from homodimers of the core protein (Cp, Fig. 1A). These dimers predominantly form *T* = 4 icosahedral capsids composed of 240 subunits, with *T* = 3 particles (180 subunits) also observed under certain conditions.^13,14^ Structural studies show that Cp adopts a largely *α*-helical fold and dimerizes through an intradimer interface that supports the characteristic capsid spike.^14–16^ Capsid assembly proceeds through a nucleation-dependent pathway, beginning with the formation of small oligomeric intermediates, often described as hexamers or trimers-of-dimers (Fig. S1A), which provide a curved scaffold for subsequent growth.^17–19^ Beyond serving as the structural building blocks of the capsid, Cp dimers participate in multiple functional transitions, including packaging of the pregenomic RNA/polymerase complex, reverse transcription within maturing nucleocapsids, intracellular trafficking, and envelopment during virion morphogenesis.^9–11,16^ These processes require a balance between cooperative protein–protein interactions and conformational adaptability, making capsid behavior sensitive to local changes in intersubunit contact geometry and stability.^10,18^

**Figure 1:**
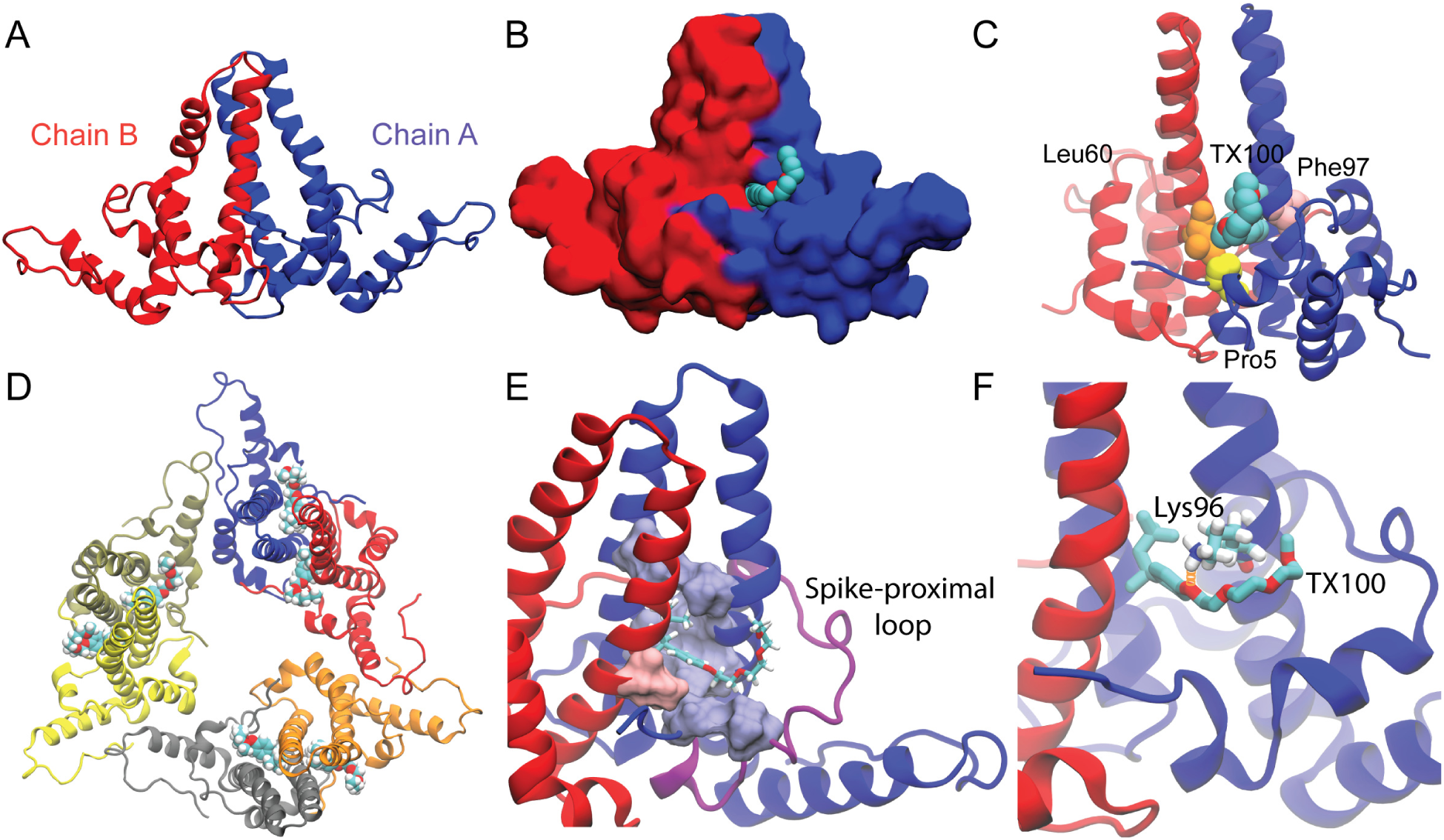
Structural context and TX100 interactions at the HBV core protein (Cp) intradimer pocket. **(A)** Side view of the Cp dimer, with chains A and B in blue and red, respectively. **(B)** Surface representation of the Cp dimer with TX100 bound at the intradimer pocket. Hydrogen atoms are omitted. **(C)** Cartoon representation showing envelopment- or secretion-associated residues: chain B Leu60 in orange, chain A Pro5 in yellow, and chain A Phe97 in pink. **(D)** TX100-bound Cp149 hexamer simulation model with one ligand in each of the six intradimer pockets. **(E)** Residues forming persistent hydrophobic contacts with TX100 (*>*80% occupancy): chain A Pro5, Val13, Ala58, Leu65, Val93, Phe97, and Leu100, and chain B Leu60. The spike-proximal loop (residues 13–30) is shown in purple. **(F)** Representative hydrogen bond between TX100 and Lys96 at the pocket entrance.

Cp has been validated as a druggable antiviral target by capsid assembly modulators (CAMs), which bind a well-characterized site at the interdimer interface (Fig. S1B) and alter capsid assembly kinetics, morphology, or stability.^20–26^ These studies demonstrate that small molecules can modulate HBV capsid behavior by engaging Cp protein–protein interfaces, motivating exploration of additional functionally relevant Cp sites.^11,12^ Beyond the canonical interdimer CAM site, a second Cp site has emerged that is structurally distinct and functionally linked to virion secretion: a hydrophobic pocket at the intradimer interface near the base of the capsid spike (Fig. 1B).^27,28^ Cryo-EM analysis identified bound Triton X-100 (TX100) molecules in this pocket and revealed localized structural responses, most notably a rotameric change of Phe97, without large backbone rearrangement.^28^ Complementary solid-state NMR studies further reported pocket-proximal chemical shift perturbations and dynamics consistent with a binding-coupled local conformational transition, supporting the existence of a TX100-responsive intradimer site.^27,29^ Systematic modification of TX100 analogs showed that binding depends strongly on hydrophobic and aromatic features, while tolerating variation in the hydrophilic headgroup, indicating a chemically selective and potentially ligandable pocket.^30^

Importantly, mutations in the structural neighborhood of this intradimer pocket are associated with pronounced secretion phenotypes (Fig. 1C). Naturally occurring Cp substitutions such as P5T and L60V reduce virion secretion while largely preserving intracellular replication.^31^ In contrast, the F97L substitution produces an immature secretion phenotype, with secreted particles enriched in incomplete viral DNA, implicating this region in coordinating nucleocapsid maturation with envelopment and release.^32,33^ Independent mutational mapping further identified residues near the base of the spike, including Leu60 and Lys96, as critical for efficient virion production.^34^ Collectively, these observations highlight the intradimer interface as a functionally sensitive region and motivate its characterization as a secretion-linked, ligandable pocket on Cp.

While cryo-EM and solid-state NMR have established pocket occupancy and local structural responses, the conformational ensemble and interaction preferences of this intradimer site remain incompletely defined. Atomistic molecular dynamics (MD) simulations provide a complementary approach to characterize pocket flexibility and the persistence of local protein–ligand interactions that are averaged in static structures. Previous simulations of HBV Cp oligomers and capsids have demonstrated that Cp dynamics and local structural variability are closely linked to function and ligand response.^35–41^ In this context, MD simulations enable analysis of how ligand binding is accommodated within the pocket and whether interactions remain localized or extend to adjacent structural regions.

To convert insights from structural dynamics into testable chemical hypotheses, fragment-based mapping can be combined with MD sampling. Conventional molecular docking is widely used for structure-based virtual screening and can efficiently generate candidate binding poses and rank ligands by approximate scoring functions.^42–45^ However, docking-based prioritization can be sensitive to the selected receptor conformation and to approximate treatment of protein flexibility, solvation, and interaction energetics, which can be particularly limiting for shallow or conformationally heterogeneous pockets. The Site-Identification by Ligand Competitive Saturation (SILCS) method addresses these challenges by generating three-dimensional functional group free-energy maps (FragMaps) from ensemble-based fragment sampling, thereby identifying favorable locations for hydrophobic, aromatic, and hydrogen-bonding interactions.^46^ Because these maps are derived from ensemble sampling, SILCS is particularly well suited for characterizing pockets that are shallow, conformationally heterogeneous, or only transiently formed, features that are consistent with the intradimer site described above. FragMap-derived pharmacophores can then be integrated with virtual screening workflows to prioritize ligands based on the presence and spatial arrangement of interaction features, enabling feature-based selection beyond docking energetics alone.^47^

Here, we integrate atomistic MD simulations, SILCS-based fragment mapping, feature-based virtual screening, and complementary binding assays to develop a workflow for evaluating ligand candidates at dynamic protein-interface pockets. In the HBV Cp intradimer pocket, MD simulations of TX100-bound Cp define a reference binding mode and identify interaction features that may support ligand engagement beyond the canonical hydrophobic cavity. These features are encoded in a SILCS-derived pharmacophore model and used to screen large compound libraries based on spatial interaction patterns rather than docking scores alone. Candidate ligands are then evaluated by saturation-transfer difference nuclear magnetic resonance (STD NMR) and surface plasmon resonance (SPR) to assess ligand–Cp interaction, followed by MD simulations to determine whether proposed pocket-bound poses are dynamically maintained and whether interactions remain localized or extend toward adjacent structural regions. Applied to the HBV Cp intradimer pocket, this workflow identifies compound B3 as a ligand candidate with experimental support and an MD-supported extended binding mode. More broadly, this study demonstrates how dynamic fragment mapping and multi-assay triage can define tractable ligand-discovery strategies for heterogeneous protein-interface pockets.

## 2 Results

### 2.1 Molecular Dynamics Analysis of TX100 Binding Reveals Key Interactions in the HBV Capsid Intradimer Pocket

To guide ligand discovery, we established a reference model of the intradimer pocket in a ligand-bound state using TX100, which has been experimentally observed to bind this site. The TX100-bound Cp dimer from the cryo-EM structure (PDB ID: 7PZK) was aligned to a closed-form Cp149 hexamer scaffold derived from PDB ID 4BMG, and ligand coordinates were transferred to generate a symmetrically occupied hexamer with one TX100 molecule per pocket (Fig. 1D). The hexamer was chosen as a computationally tractable higher-order assembly that retains the trimer-of-dimers organization and provides a more complete local intersubunit environment around the intradimer pocket than an isolated dimer.

All simulations were performed in three independent 160-ns replicas, with the first 10 ns excluded as equilibration. Trajectory stability was confirmed by root-mean-square deviation (RMSD) (Fig. S2). Hydrophobic contact analysis revealed a cluster of high-occupancy interactions defining a continuous apolar cavity within the intradimer pocket (Figs. 1E, S3A, and S4). The hydrophobic head group and phenolic ring of TX100 insert deeply into the cavity, while the polyethylene glycol tail remains largely solvent-exposed and does not form persistent contacts. The pocket interior is lined by hydrophobic residues from chain A, including Leu100, Phe97, Val93, Ala58, and Leu65, with additional contributions from Leu60 of chain B at the dimer interface. Notably, Pro5 and Val13 of chain A are located within a loop at the base of the spike (Fig. 1E), adjacent to but not part of the contiguous hydrophobic cavity. This loop forms part of the structural linkage between the intradimer interface and the spike region. Hydrophobic contacts involving these residues remain localized near the pocket entrance and do not extend further along the loop, indicating that TX100 binding is largely confined to the cavity. However, the proximity of this loop to the binding site suggests that ligand extension toward this region could provide a way to probe whether pocket binding can influence spike-associated conformational dynamics.

Complementing the hydrophobic interactions, hydrogen-bond analysis identifies a limited set of polar contacts at the intradimer pocket (Fig. S3B). Only Lys96 (Fig. 1F) and Gln99 (Fig. S5) from chain A exhibit appreciable hydrogen-bond occupancy with TX100. Lys96 acts as the primary donor, forming a stable interaction with the first ether oxygen of the ligand (∼75% occupancy), whereas Gln99 participates in more transient contacts (∼22%), primarily with the second ether oxygen along the ligand chain. These interactions are localized near the pocket entrance and involve the more solvent-exposed portion of the ligand. Overall, TX100 binding is dominated by hydrophobic interactions, with a small number of polar contacts, primarily mediated by Lys96, providing additional stabilization at the pocket periphery.

Comparison of apo and TX100-bound simulations reveals ligand-induced side-chain rearrangements at the intradimer pocket (Figs. S6 and S7). In the apo state (Fig. S7A), Phe97 is oriented toward the pocket interior, whereas in the TX100-bound state (Fig. S7B), it is displaced deeper into the cavity. In parallel, Lys96 reorients toward the pocket entrance, enabling hydrogen-bond formation with the ether oxygen atoms of TX100. Together, these observations indicate that ligand binding is accommodated through localized side-chain rearrangements that adapt to the pocket geometry.

### 2.2 SILCS-Based Mapping of Interaction Preferences and Pharmacophore Construction

To characterize the interaction preferences of the intradimer pocket, we applied SILCS.^46,47^ Guided by the ligand-induced conformational changes observed in the TX100-bound simulations, the Cp149 tetramer derived from the TX100-bound structure (PDB ID: 7PZK) was used as the starting model, with all ligand molecules removed. The protein was simulated in the presence of a diverse set of probe molecules representing key functional groups, including aliphatic, aromatic, hydrogen-bond donor, and hydrogen-bond acceptor moieties. The complete probe set is described in the Methods. The resulting trajectories were used to generate grid free energy (GFE) FragMaps, which quantify the spatial preferences of these functional groups around the intradimer pocket.

The generated FragMaps are shown in Fig. 2B as isosurfaces representing regions of favorable interactions for different functional groups. The Cp149 tetramer contains four equivalent intradimer pockets; comparison across these sites shows that apolar features (cyan isosurfaces) are largely conserved, whereas polar features exhibit localized variability (Fig. S8). For subsequent analysis, we focused on the pocket formed between chains A and B on the interior-facing side of the tetramer, which displays a more defined and contiguous distribution of polar interaction features (circled in Fig. 2B).

**Figure 2:**
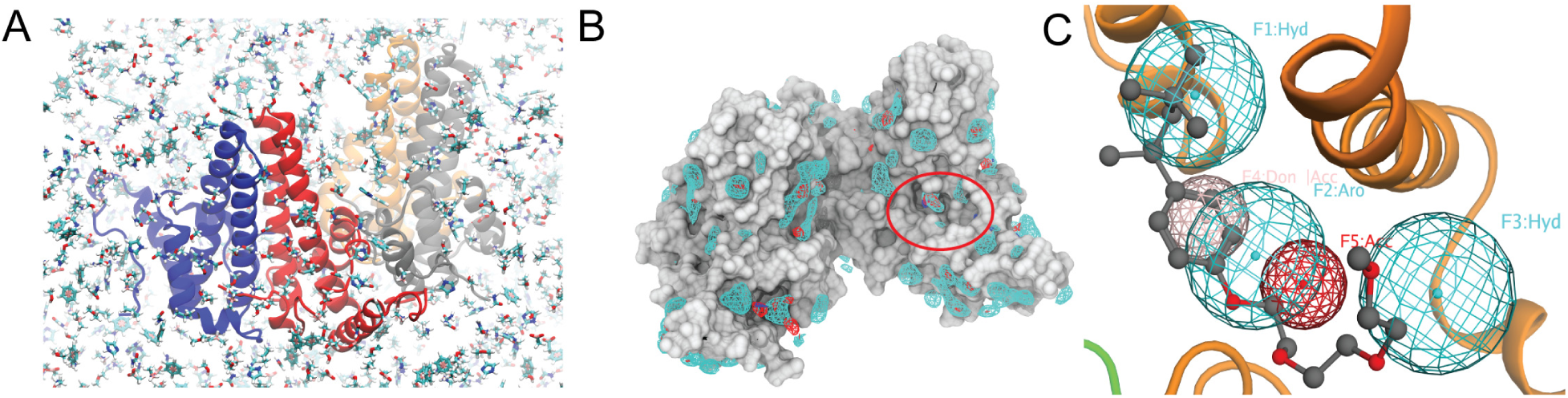
SILCS characterization and pharmacophore construction at the Cp intradimer pocket. **(A)** Representative SILCS simulation snapshot showing the Cp149 tetramer and surrounding fragment probes. **(B)** FragMaps highlighting apolar (cyan), hydrogen-bond acceptor (red), and hydrogen-bond donor (blue) regions around the intradimer pocket (circled). **(C)** Pharmacophore model overlaid with TX100 at the Cp A–B intradimer interface. Features are shown as wireframe spheres (cyan, apolar; red, acceptor; blue, donor; pink, donor/acceptor overlap). The excluded-volume constraint is omitted for clarity and is shown with the final five-feature model in Fig. S10A.

To define discrete interaction sites, FragMap regions within the selected pocket were clustered based on spatial proximity and free energy favorability.^47^ This process yielded eight candidate features (Fig. S9), from which five representative pharmacophore features were selected (Fig. 2C) based on spatial coherence and consistency with the ligand binding geometry observed in the TX100-bound simulations. In Fig. 2C, features 1–3 define apolar regions that collectively form the hydrophobic core of the pocket, with their spatial overlap relative to the TX100 binding pose. Feature 1 overlaps with the position of the aliphatic head group of TX100, while Feature 2 corresponds to the phenolic ring region near the cavity entrance. Feature 3 extends toward the loop at the base of the spike and was included to capture a hydrophobic extension site adjacent to Pro5 and Val13. This region is not engaged by TX100 and provides a potential route for ligand interactions beyond the canonical hydrophobic cavity. Feature 4 defines a hydrogen-bond donor site near Glu57, representing a potential polar interaction that is not reflected in the observed TX100 binding mode. Feature 5 defines a hydrogen-bond acceptor site in proximity to Lys96 and Gln99. This feature lies close to, but does not perfectly overlap with, the first ether oxygen of TX100, consistent with the hydrogen-bonding interactions observed in the TX100-bound simulations while indicating a broader favorable region for acceptor placement. Together, these features define a spatially connected set of favorable interaction regions from the pocket interior toward the spike-proximal loop, providing a structural basis for selecting ligands with the potential to extend beyond the canonical hydrophobic cavity.

### 2.3 Feature-Based Screening Prioritizes Eight Compounds for Experimental Evaluation

Based on the SILCS-derived pharmacophores, we implemented a sequential triage workflow to identify and evaluate compounds compatible with the intradimer pocket. The workflow combined pharmacophore matching, physicochemical filtering, docking-pose evaluation, experimental binding assessment, and MD-based characterization of pose stability and spatial extent. In the initial pharmacophore-screening stage, the three apolar features were treated as essential to define the ligand scaffold, while the two polar features were optional, with at least one required to be satisfied. Feature-based screening was carried out using the Pharmacophore Search module in MOE.^48^ We screened approximately 4.7 million compounds from the Enamine Screening Collection,^49^ using precomputed ligand conformations stored in molecular database (MDB) format. Compounds were evaluated for geometric compatibility with the pharmacophore and exclusion volume constraints using these precomputed conformations (Fig. S11). This approach avoids explicit energy evaluation during the initial screen and enables efficient evaluation of large compound libraries, yielding ∼78,000 candidates that satisfied the pharmacophore criteria.

The pharmacophore-derived candidates were then filtered by predefined physicochemical criteria and reactive-group exclusions. Molecular descriptors were calculated using the Descriptor Calculator in MOE.^48^ The filters included constraints on molecular size, hydrogen-bonding capacity, flexibility, polarity, lipophilicity, and molar refractivity. Compounds containing predefined reactive functional groups were also excluded. The cutoff values used for physicochemical filtering are listed in Table S1. This step reduced the candidate set to approximately 10,000 compounds for molecular docking.

The filtered compounds were subjected to docking to evaluate their compatibility with the intradimer pocket. While pharmacophore screening selects compounds based on the presence and spatial arrangement of key interaction features, docking provides an additional assessment of steric and energetic fit within the binding site. Docking was performed using the MOE workflow, initialized from pharmacophore-matched poses and allowing local optimization with induced-fit refinement. Compounds with docking scores better than −8 kcal/mol were retained, reducing the candidate set to 120 molecules (Figs. S12–S16). Docking scores and selected physicochemical properties of the compounds chosen for experimental evaluation are summarized in Table S2.

From the 120 compounds retained after docking-score filtering, docking poses were visually inspected to confirm preservation of the pharmacophore-matched geometry after induced-fit refinement, including occupancy of the hydrophobic intradimer pocket, satisfaction of the required apolar features and at least one polar feature, absence of major steric clashes, and appropriate placement of the feature-matched functional group. Compounds were then selected to capture differences in scaffold, ligand orientation, spatial extent, and predicted residue contacts, while closely related structures, highly similar interaction patterns, and ligands containing large regions not involved in the proposed pocket interactions were deprioritized. Commercial availability was also required. Together, these criteria reduced the 120 docking-retained compounds to eight candidates for experimental evaluation (Fig. 3). The A-series was selected to satisfy Feature 4 near Glu57, the B-series was selected to satisfy Feature 5 near Lys96 and Gln99, and B4 satisfied both polar features.

**Figure 3:**
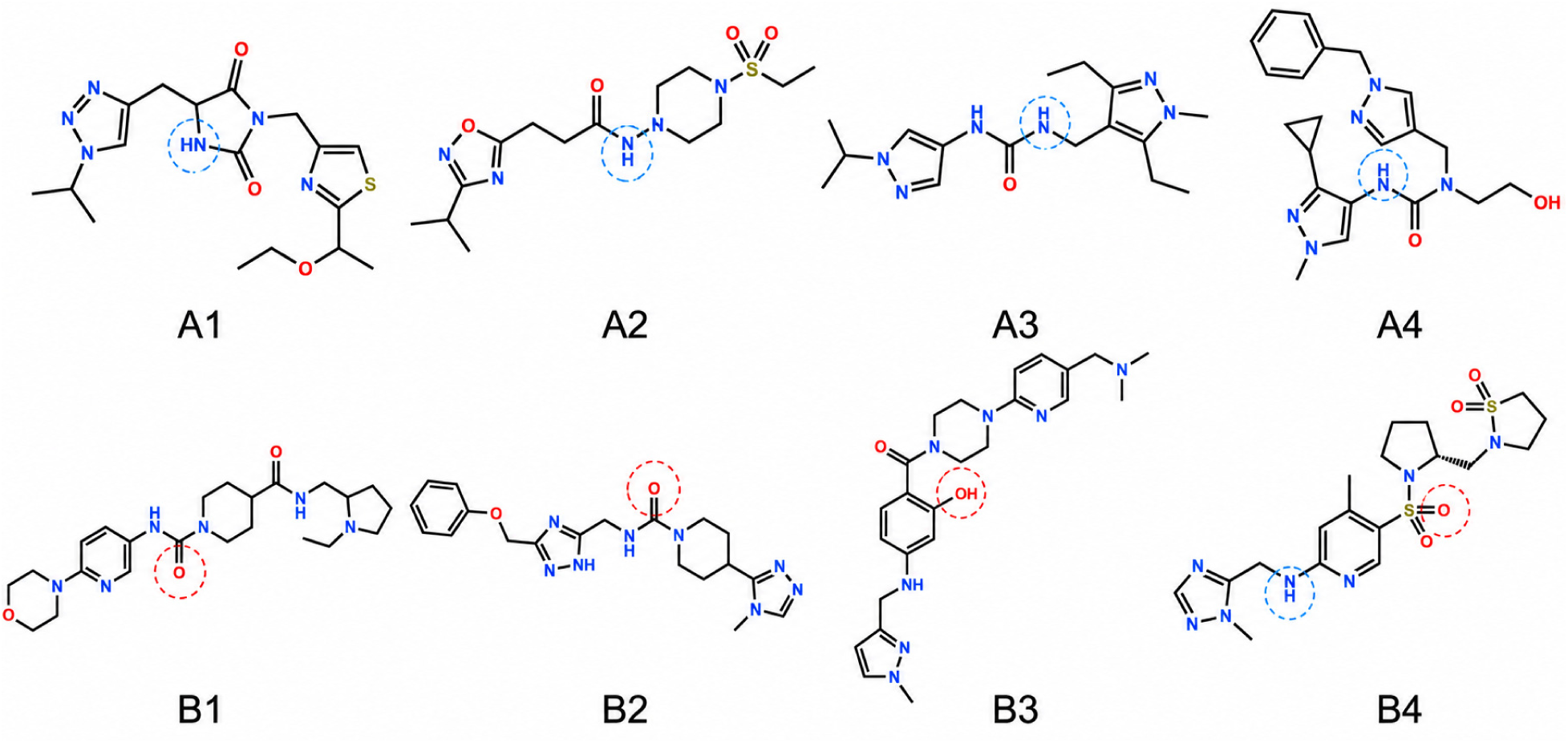
Structures of the eight compounds selected for experimental evaluation based on pharmacophore-guided virtual screening. Blue circles indicate ligand groups satisfying Feature 4, corresponding to the hydrogen-bond donor feature near Glu57. Red circles indicate ligand groups satisfying Feature 5, corresponding to the hydrogen-bond acceptor feature near Lys96 and Gln99. For B4, both Feature 4 and Feature 5 are satisfied.

### 2.4 Experimental Binding Assays Distinguish B3 from Other Candidate Ligands

Experimental characterization of the selected compounds was performed using STD NMR and SPR. STD NMR detects ligand–protein interactions through saturation transfer from Cp149 to bound ligands in solution,^50^ and the distribution of STD signals across ligand resonances can provide qualitative information about ligand contact patterns. SPR provides an orthogonal measurement of concentration-dependent ligand binding to immobilized HBcAg-biotin and enables estimation of apparent binding affinity.

STD NMR experiments were first used to assess ligand interaction with Cp149 in solution. STD responses were evaluated by comparing ligand–Cp149 samples with ligand-only controls acquired under identical conditions. Among the eight compounds tested, A1 (Fig. S17) and A2 (Fig. S18) showed no obvious STD signals above the corresponding controls, whereas A3 (Fig. S19) and B1 (Fig. S21) exhibited weak or ambiguous responses due to low signal intensity. Compounds A4 (Fig. S20), B2 (Fig. S22), B3 (Fig. S23), and B4 (Fig. S24) showed detectable STD signals above the ligand-only controls, consistent with interaction with Cp149. The distribution of STD signals also suggested differences in ligand contact patterns: B2 and B4 showed more localized responses, whereas A4 and B3 displayed signals across multiple ligand resonances, consistent with more distributed ligand–protein contacts. Although STD NMR can provide information about ligand binding epitopes and orientation, it does not by itself uniquely identify the protein binding site. These observations were therefore interpreted as qualitative evidence for ligand–Cp149 interaction and differences in contact pattern.

SPR was next used to evaluate concentration-dependent binding of the eight compounds to immobilized HBcAg-biotin. Under the conditions tested, A2, A3, A4, B2, and B4 showed no appreciable SPR response (Fig. S25). A1 and B1 produced weak responses, with estimated affinities exceeding 1 mM. In contrast, B3 showed the clearest concentration-dependent response and the strongest measurable affinity among the compounds tested, yielding an estimated *K*_D_ of 563.2 ± 93.0 *µ*M (Fig. 4C and Fig. S25). B3 was therefore prioritized for further structural characterization, with its detectable STD NMR signals providing independent support for interaction with Cp149.

**Figure 4:**
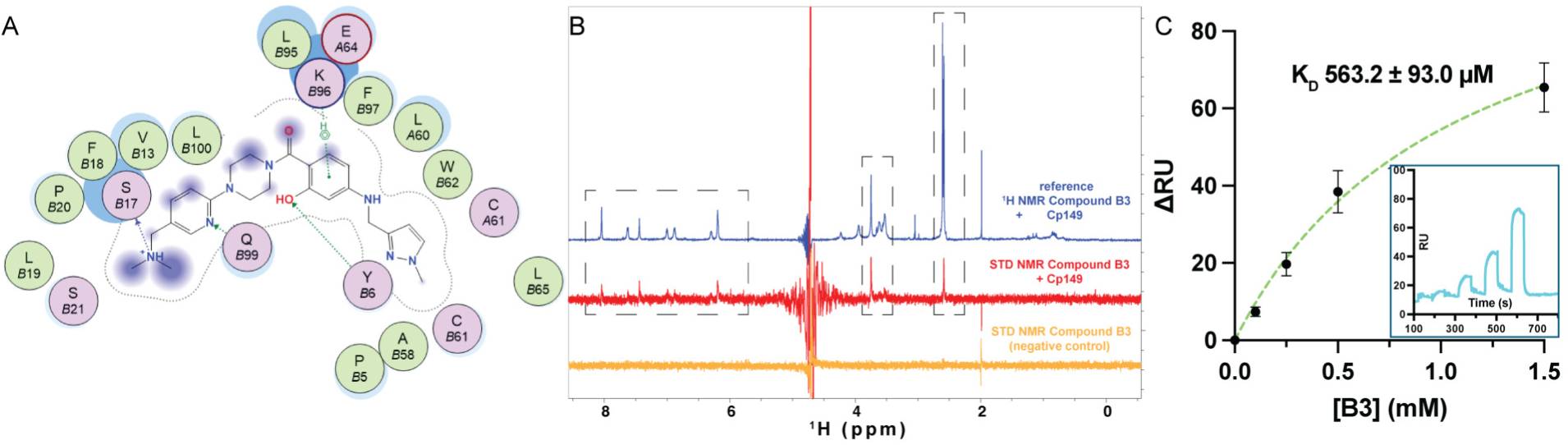
Experimental and computational characterization of compound B3. **(A)** Representative two-dimensional interaction diagram of B3 within the Cp149 intradimer pocket. Hydrogen bonds, hydrophobic contacts, and surrounding pocket residues are shown. See also Fig. S26. **(B)** STD NMR spectra of B3 in the presence of Cp149. The blue spectrum is the one-dimensional ^1^H NMR spectrum of B3 in the presence of Cp149, the red is the corresponding STD spectrum, and the orange is the ligand-only negative control acquired under identical conditions. Brackets indicate STD signals absent from the control. **(C)** SPR analysis of B3 binding to immobilized HBcAg-biotin. The main panel shows the steady-state binding response as a function of B3 concentration with the fitted equilibrium dissociation constant (*K*_D_). Error bars represent the standard deviation from three independent experiments. The inset shows a representative SPR sensorgram obtained at increasing B3 concentrations (0.1, 0.25, 0.5, and 1.5 mM).

The differences between the STD NMR and SPR results likely reflect the distinct physical basis and experimental format of the two assays. STD NMR can detect weak, transient, or heterogeneous ligand–protein contacts in solution, whereas SPR requires a concentration-dependent response from binding to immobilized HBcAg-biotin under flow conditions. Thus, STD-positive but SPR-negative compounds may form solution-state contacts that are too weak, transient, heterogeneous, or assay-condition dependent to generate a measurable SPR response. Conversely, the weak SPR responses observed for A1 and B1 may reflect low-affinity interactions that were less clearly resolved by STD NMR.

B3 showed STD NMR signals across multiple ligand resonances (Fig. 4B), suggesting a more distributed pattern of ligand–protein contact than compounds with localized STD responses. Its proposed binding pose places B3 within the intradimer pocket while extending toward the spike-proximal loop region (Fig. 4A), in contrast to the TX100 reference mode, which is dominated by hydrophobic occupancy of the core cavity. This proposed pose motivated subsequent MD analysis to determine whether the extended binding mode and associated contacts were dynamically maintained.

### 2.5 MD Simulations Support an Extended Intradimer-Pocket Binding Mode for B3

To assess whether the proposed intradimer-pocket binding poses could be maintained dynamically, MD simulations were performed for the eight selected ligands using docking-derived starting poses in the Cp149 hexamer. For each protein-ligand system, a single ligand was placed in the outward-facing intradimer pocket formed by chains C and D. For each system, three independent simulations of 160 ns were performed, and the first 10 ns of each trajectory were excluded as equilibration. These simulations were used to evaluate binding-mode stability and contact patterns, but not to directly determine binding affinity or thermodynamic stability.

For A1, its instability contrasts with the weak SPR response, suggesting that its low-affinity SPR signal may arise from a binding mode not captured by the simulated intradimer-pocket pose or from weak, nonspecific, or heterogeneous interactions with HBcAg. In contrast, A3, B1, B2, and B4 maintained localized pocket-bound configurations, in which one hydrophobic group remained buried in the intradimer cavity while the rest of the ligand was largely solvent-exposed (Fig. S27A,C,D,F). However, these localized MD poses were not consistently supported by both STD NMR and SPR, indicating that maintenance of a local pose over the 150-ns analysis window is not sufficient to establish robust experimental binding. Among the simulated compounds, B3 was the only ligand that maintained an extended intradimer-pocket pose across independent simulations, consistent with its prioritization based on the SPR results.

B3 displayed a more extended binding mode compared with TX100 and the other candidate ligands. Across independent simulations, B3 maintained a broader intradimer-pocket binding pose in which one portion of the ligand remained anchored within the hydrophobic cavity while the rest of the molecule extended toward the spike-proximal loop region (Fig. 5A and Fig. S27E). This behavior contrasts with the TX100 reference mode, where ligand engagement is dominated by hydrophobic occupancy of the core cavity and remains largely confined near the pocket entrance. It also differs from the localized poses observed for A3, B1, B2, and B4, in which pocket occupancy was maintained but extended contacts with adjacent structural elements were limited.

**Figure 5:**
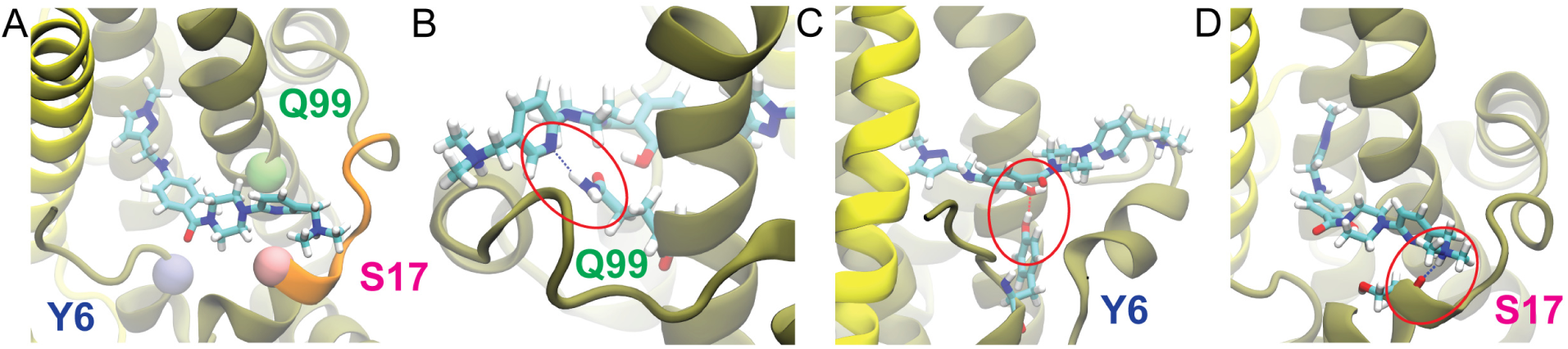
Binding mode of ligand B3 from MD simulations. **(A)** Representative binding pose of B3, with chain C shown in yellow and chain D in tan. The C*α* positions of residues involved in hydrogen-bonding interactions are indicated by spheres (Gln99 in lime, Tyr6 in iceblue, and Ser17 in pink). The loop region of chain D (residues 18–22) is highlighted in orange. **(B–D)** Zoomed-in views of hydrogen-bonding interactions with surrounding residues: (B) Gln99, (C) Tyr6, and (D) the backbone of Ser17.

The extended B3 pose was stabilized by a set of polar interactions distributed around the ligand. The nitrogen atom within the pyridine ring formed a hydrogen bond with the side chain of Gln99 near the pocket entrance, with an occupancy of 37% across the analyzed trajectories (Fig. 5B). The hydroxyl group on the central phenyl ring formed a hydrogen bond with Tyr6, with an occupancy of 84% (Fig. 5C). In addition, the terminal tertiary amine, predicted to be protonated at physiological pH, interacted with the backbone carbonyl of Ser17 in the spike-proximal loop region, with an occupancy of 67% (Fig. 5D). These interactions anchor B3 from multiple directions and provide a structural basis for maintaining contact between the hydrophobic intradimer pocket and the adjacent loop region.

In addition to maintaining extended contacts outside the core cavity, B3 binding was also associated with local rearrangement of the intradimer pocket. Because TX100 binding is associated with displacement of Phe97 from the apo-like pocket-occluding orientation, we examined whether B3 produced a similar side-chain response. Analysis of Phe97 conformations showed that B3 binding also shifted Phe97 away from the apo-like orientation, consistent with local pocket adaptation upon ligand binding (Fig. S28). This behavior supports the similarity between B3 and TX100 in hydrophobic pocket engagement, while the additional contacts formed by B3 distinguish it from the more localized TX100 binding mode.

Together, the simulations support a model in which B3 engages the intradimer pocket through both hydrophobic anchoring and extended contacts outside the canonical cavity. This binding mode provides a plausible structural explanation for the broader STD NMR signal distribution observed for B3 and is consistent with its measurable SPR response. Although MD simulations do not independently establish binding affinity, they support an extended binding-mode hypothesis for B3 and provide a structural basis for further optimization.

## 3 Discussion

The intradimer interface pocket of the HBV core protein represents a structurally distinct and functionally relevant site for small-molecule modulation. Unlike the interdimer CAM-binding site, this intradimer pocket is less extensively characterized and lies within a conformationally heterogeneous protein–protein interface, making its interaction preferences and ligandability less straightforward to define using conventional structure-based approaches. Previous studies of TX100 and related compounds established that the site can accommodate hydrophobic ligands, but these ligands primarily occupy the core cavity with limited engagement of adjacent structural regions.^27,30^ Whether interactions at this site can be extended beyond the canonical hydrophobic pocket, and how such ligands might be identified, has remained unclear.

A central outcome of this work is the application of a feature-based screening strategy to the HBV Cp intradimer pocket, rather than relying on docking-score-driven selection alone. Conventional docking is useful for generating poses and assessing steric compatibility, but docking scores can be sensitive to receptor conformation, scoring function, and the treatment of protein flexibility and solvation. These limitations are particularly relevant for dynamic protein-interface pockets, where productive engagement may depend on satisfying a specific spatial arrangement of interactions rather than simply occupying the cavity. Here, SILCS-derived pharmacophores built from fragment free-energy maps sampled in an explicitly solvated, dynamic protein environment were combined with physicochemical and reactive-group filtering, docking-pose evaluation, and subsequent experimental and computational characterization. Together, these steps narrowed the initial library of approximately 4.7 million compounds to eight candidates for experimental evaluation.

Of the eight candidates, B3 was prioritized because it showed the clearest concentration-dependent SPR response and the strongest measurable affinity among the compounds tested. Its detectable STD NMR signals provided independent support for interaction with Cp149, while MD simulations suggested an extended binding mode in which B3 remained anchored within the intradimer pocket and extended toward the spike-proximal loop region. This pose was associated with contacts near the pocket entrance and with residues in the adjacent loop region, thereby bridging the canonical hydrophobic cavity and nearby structural elements. The resulting model provides a structural rationale for the broader STD NMR signal distribution observed for B3 and illustrates how ligand engagement at this site may extend beyond the localized binding mode observed for TX100.

These findings extend prior studies of the intradimer pocket by placing ligand engagement at this site in a broader structural context. TX100 and related compounds established that the site is ligandable, but their reported binding modes are dominated by hydrophobic occupancy of the core cavity.^30^ For further development toward drug-like ligands, greater incorporation of polar functionality may be desirable to improve aqueous solubility and provide additional directional interactions beyond the hydrophobic core. Consistent with this design principle, the B3 binding model suggests that ligand engagement need not remain confined to the hydrophobic cavity when polar and apolar interaction features are appropriately organized. This distinction is relevant because the adjacent spike-proximal region contains residues implicated in capsid maturation, envelopment, and virion secretion. Although the present study does not directly test functional consequences, the B3 model provides a basis for optimizing ligands that couple intradimer-pocket occupancy with engagement of adjacent structural elements.

Several limitations should be noted. The experimental and computational readouts were not fully concordant for all compounds, reflecting the challenges of characterizing weak ligands at a dynamic protein-interface pocket. STD NMR can detect weak or transient ligand–protein contacts in solution and provide qualitative information about the ligand binding pose, but does not necessarily identify the binding site; SPR provides concentration-dependent binding information but may be influenced by immobilization and assay conditions; and MD simulations depend on the starting pose, force field, and sampling. To mitigate the limitations of any individual method, we combined orthogonal solution- and surface-based binding assays with MD-based assessment of pose stability and contact patterns. Concordance across these independent readouts therefore provides stronger support for ligand interaction and the proposed binding-mode hypotheses than any single measurement alone. Accordingly, the proposed binding modes should be interpreted as MD-supported structural models rather than definitive assignments. In addition, this study focuses on structural and biophysical characterization rather than direct assessment of capsid assembly, secretion, or viral replication. Given the weak affinities of the current compounds, biological testing will likely be most informative after further optimization of affinity, solubility, and binding-mode stability.

Overall, this work establishes a feature-based screening strategy for ligand discovery at the conformationally heterogeneous HBV Cp intradimer interface. Integration of SILCS fragment mapping, pharmacophore-guided screening, biophysical assays, and MD simulations identified B3 as a ligand candidate with experimental support and a binding mode extending from the hydrophobic pocket toward the spike-proximal loop. More broadly, the study illustrates how interaction-feature-driven screening can complement conventional docking for challenging protein-interface pockets.

## 4 Methods

### 4.1 MD simulations

All MD simulations were performed using NAMD3.^51^ The CHARMM36m force field was used for protein representation, with CHARMM36 and TIP3P parameters applied to ions and water, respectively.^52–54^ Simulations of the HBV Cp149 hexamer were based on the crystal structure of the closed conformation (PDB ID: 4BMG).^55^ For the TX100-bound system, a Cp dimer extracted from the TX100-bound structure (PDB ID: 7PZK) was aligned to the Cp149 hexamer scaffold, and the ligand coordinates were transferred to each intradimer interface to generate a symmetrically occupied hexamer. The ligand molecules were parameterized using the CGenFF force field.^56^

All systems were solvated in explicit water under periodic boundary conditions and ionized to a physiological salt concentration of 0.15 M NaCl. Prior to production simulations, systems were energy minimized and equilibrated in three stages. First, the protein and ligand were positionally restrained while the system was minimized for 2,000 steps and equilibrated for 1 ns. Second, restraints were reduced to the protein backbone, followed by 2,000 steps of minimization and 1 ns of equilibration. Finally, all restraints were released, and the system was equilibrated for an additional 1 ns before production simulations. Temperature was maintained at 310 K using Langevin dynamics, and pressure was maintained at 1 atm using a Langevin piston barostat. Covalent bonds involving hydrogen atoms were constrained, allowing a 4-fs integration time step.^57^ Short-range nonbonded interactions were computed with a cutoff of 12 Å and a switching function applied between 10 and 12 Å, while long-range electrostatics were treated using the particle-mesh Ewald method.^58^

For each system, three independent simulations were performed, each for 160 ns. The first 10 ns of each trajectory were discarded as equilibration, and the remaining trajectories were used for analysis. Trajectory visualization was performed using VMD.^59^

### 4.2 Analysis of MD Trajectories

All trajectory analyses were performed using MDAnalysis.^60^ Protein structural stability was assessed by calculating the RMSD of C_α_ atoms relative to the initial frame of each trajectory (frame 0). Before RMSD calculations, each trajectory frame was aligned to the initial frame using the protein C_α_ atoms. RMSD values were computed for all frames and averaged across replicas. For the TX100-bound system, ligand RMSD was additionally calculated using heavy atoms of the ligand.

Hydrophobic interactions between protein residues and TX100 were quantified based on a distance cutoff of 4.5 Å between non-hydrogen atoms. Hydrophobic residues were defined as Ala, Val, Leu, Ile, Met, Phe, Trp, Tyr, and Pro. A contact was considered present in a given frame if any heavy atom of a residue was within the cutoff distance of ligand heavy atoms. Contact occupancy was calculated as the fraction of analyzed frames in which the contact was present. For symmetry-related binding sites, interactions were mapped onto equivalent positions and averaged across ligands and replicas to obtain symmetry-collapsed occupancies.

Hydrogen bonds between protein residues and TX100 were identified using geometric criteria with a donor–acceptor distance cutoff of 3.5 Å and a donor–hydrogen–acceptor angle cutoff of 130^◦^. Only protein-to-ligand hydrogen bonds were considered, with protein atoms acting as donors and ligand oxygen atoms as acceptors. Hydrogen-bond occupancy was computed as the fraction of analyzed frames in which a given interaction was present, and values were averaged across symmetry-related sites and independent replicas. For B3, hydrogen-bond occupancies supporting the proposed extended binding mode were calculated using the same geometric criteria and averaged across the three independent replicas.

Side-chain conformational changes were characterized by analyzing dihedral-angle distributions (*χ* angles) for residues throughout the protein. Dihedral angles were computed for all relevant residues and binned into histograms with a bin width of 15^◦^. Differences between apo and TX100-bound simulations were quantified using a circular Wasserstein distance metric, which measures redistribution of angular populations while accounting for periodicity. Residues exhibiting significant rotamer redistribution were identified using a threshold of 30^◦^, and results were averaged across replicas after combining symmetry-related chains.

#### SILCS Simulations and Pharmacophore Modeling

SILCS simulations were performed to characterize functional group interaction preferences of the intradimer pocket.^47^ The Cp149 tetramer derived from the TX100-bound structure (PDB ID: 7PZK) was used as the starting model, with all ligand molecules removed while retaining the protein coordinates. The simulations were carried out using an explicit-solvent MD framework in which the protein is immersed in an aqueous solution containing a mixture of small organic probe molecules representing common functional groups. The probe set included small organic fragments representing key functional groups, specifically propane (aliphatic), benzene (aromatic), methanol and formamide (neutral hydrogen-bond donor and acceptor), acetaldehyde (hydrogen-bond acceptor), methylammonium (positively charged donor), and acetate (negatively charged acceptor), consistent with the extended SILCS-Pharm protocol.^47^ During the simulations, probe molecules and water compete for interaction with the protein surface, enabling identification of energetically favorable binding regions while accounting for protein flexibility and desolvation effects.^46,47^

Ten independent SILCS simulations of 50 ns each were performed with randomly distributed probe molecules at comparable concentrations (∼0.25 M for each probe type) to enhance sampling and ensure convergence of the resulting FragMaps. Fragment–fragment aggregation was mitigated through the use of a repulsive interaction potential between probe molecules, ensuring effective sampling of protein–fragment interactions without perturbing protein–ligand interactions.^46^ To maintain structural integrity while preserving local flexibility, weak positional restraints were applied to protein C_α_ atoms with a force constant of approximately 0.1 kcal/mol/Å^2^, following standard SILCS protocols.

Trajectory data from all SILCS simulations were combined to generate FragMaps by binning the spatial distributions of probe atoms onto a grid with 1 Å spacing encompassing the protein. These probability distributions were normalized relative to bulk solvent distributions and subsequently converted into GFE maps via Boltzmann transformation, yielding quantitative representations of functional group binding preferences. Convergence of FragMaps was assessed by comparing independent subsets of trajectories using overlap metrics as described previously.^47^

Pharmacophore features were derived from the resulting GFE FragMaps using the SILCS-Pharm protocol.^47^ Voxels within the intradimer pocket exhibiting favorable GFE values were selected using default energy-based thresholds implemented in the SILCS workflow. Specifically, voxels with GFE values ≤ −1.2 kcal/mol were selected for apolar features, while thresholds of ≤ −1.0 kcal/mol were applied for hydrogen-bond donor and acceptor features. For charged features, more stringent cutoffs of ≤ −1.8 kcal/mol were used for methylammonium (donor) and acetate (acceptor) FragMaps. Selected voxels were then clustered based on spatial proximity to define FragMap features and converted into pharmacophore features, including apolar, hydrogen-bond donor, and hydrogen-bond acceptor types. Final pharmacophore models were constructed by selecting and combining features according to spatial continuity and consistency with ligand-binding geometries observed in MD simulations.

### 4.3 Pharmacophore-Based Virtual Screening

Virtual screening was performed using the pharmacophore model derived from SILCS FragMaps to identify compounds compatible with the intradimer pocket. Three apolar features were defined as essential and required to be satisfied by all candidate compounds, while two polar features were treated as optional, with at least one required to be matched. This assignment was based on the spatial distribution of favorable GFE regions and their correspondence to the TX100 binding mode, rather than on GFE magnitude alone. Feature-based screening was carried out using the Pharmacophore Search module implemented in the Molecular Operating Environment (MOE).^48^ Compounds were evaluated based on the presence and geometric compatibility of pharmacophore features in conjunction with exclusion volume constraints. Precomputed ligand conformations stored in MDB format were used, allowing efficient evaluation of feature matching without on-the-fly conformational sampling. For each compound, only the highest-scoring matching conformation was retained. The compound library screened consisted of the Enamine Screening Collection,^49^ comprising approximately 4.7 million compounds. Pharmacophore matching yielded an initial set of candidate compounds satisfying the defined feature constraints.

These candidates were subsequently filtered using predefined physicochemical criteria and reactive-group exclusions to prioritize compounds with suitable properties for subsequent evaluation. Molecular descriptors were estimated using the Descriptor Calculator in MOE.^48^ Filtering criteria included constraints on molecular size, hydrogen-bonding capacity, molecular flexibility, polarity, lipophilicity, and molar refractivity, together with exclusion of predefined reactive functional groups. Detailed cutoff values for the physicochemical filtering criteria are provided in Table S1.

### 4.4 Molecular Docking

Molecular docking was performed using the docking workflow implemented in the MOE^48^ to evaluate the compatibility of filtered compounds with the intradimer pocket. Initial ligand conformations were taken from the pharmacophore-matched poses obtained during the virtual screening stage. Docking refinement was carried out using an induced-fit protocol, allowing local optimization of both ligand geometry and protein side-chain conformations within the binding site. Candidate poses were evaluated using the MOE scoring function to estimate binding energies. Compounds with docking scores better than −8 kcal/mol were retained for further analysis.

### 4.5 STD NMR Spectroscopy

STD NMR experiments were performed on a Bruker AVIII-HD 700 MHz spectrometer at the Georgia Tech NMR Center, using pulse sequence `STDDIFFESGP.3`. Samples contained 20 µM protein and 500 µM ligand, corresponding to a 25-fold ligand excess, in 0.2× PBS buffer prepared in 90% H_2_O and 10% D_2_O. Control samples containing ligand only under identical conditions were also prepared. STD NMR spectra were acquired using selective saturation of protein resonances, with an on-resonance frequency set to −0.7 ppm, corresponding to protein methyl groups, and an off-resonance frequency set to 40 ppm. A total saturation time of 2 s was applied. Spectra were collected with 64 scans.

STD difference spectra were obtained by subtracting off-resonance spectra from on-resonance spectra, yielding signals corresponding to ligand protons affected by saturation transfer from the protein. The absence of STD signals in ligand-only control experiments confirmed that the observed STD effects arise from protein-mediated interactions. STD signals were interpreted qualitatively to assess ligand binding, where the presence and distribution of signals across ligand protons indicate regions of interaction with the protein.^50^

### 4.6 SPR experiments

HBcAg-biotin was purchased from Creative Diagnostics (catalogue number DAG-WT6839B, UniProt ID P0C6X7). SPR experiments were performed in triplicate (*n* = 3) using a Biacore T200 instrument (Cytiva) in SPR buffer (0.2× PBS, pH 7.2, 3% DMSO-*d*_6_, 0.05% v/v Tween-20). Small molecules were prepared as 30 mM stock solutions and diluted into SPR buffer. Approximately 250–450 resonance units (RU) of HBcAg-biotin were immobilized on a streptavidin-coated sensor chip. Small molecules at concentrations of 0, 0.1, 0.25, 0.5, and 1.5 mM were injected over the HBcAg-coated surface. All experiments were performed at 25 ^◦^C with a flow rate of 50 µL min^−1^. Small molecules were injected for 60 s followed by a 180 s dissociation phase. SPR response was reference-subtracted using a control flow cell lacking immobilized protein. Equilibrium dissociation constants (*K*_D_) were determined with Biacore T200 evaluation software v3.1 (Cytiva) using a surface-bound steady-state affinity analysis assuming a 1:1 binding stoichiometry. SPR sensorgrams were prepared in GraphPad Prism v10.

## Acknowledgement

This work was supported by the National Institutes of Health (NIH; R01-AI148740). Computational resources included the Phoenix cluster, which is managed by the Partnership for an Advanced Computing Environment (PACE) at the Georgia Institute of Technology. We thank the Georgia Tech NMR Center for access to instrumentation and technical support, and acknowledge the use of a Bruker AVIII-HD 700 MHz spectrometer. We are grateful to Dr. Hongwei Wu for assistance with NMR experiments. We also thank Dr. Alain Ajamian at the Chemical Computing Group for technical support and helpful discussions regarding MOE.

## 4.7 Data Availability

The dataset can be accessed via DOI: https://doi.org/10.5281/zenodo.21959223. The repository contains SILCS FragMaps, the derived pharmacophore model, and MD simulation data used to characterize ligand interactions at the HBV core protein intradimer pocket.

## 4.8 Supporting Information

The Supporting Information is available free of charge at: xxxxx. Additional figures and analyses include TX100-bound MD characterization, SILCS FragMaps and pharmacophore construction, virtual-screening results, STD NMR and SPR measurements, and ligand binding-mode characterization.

## TOC Graphic

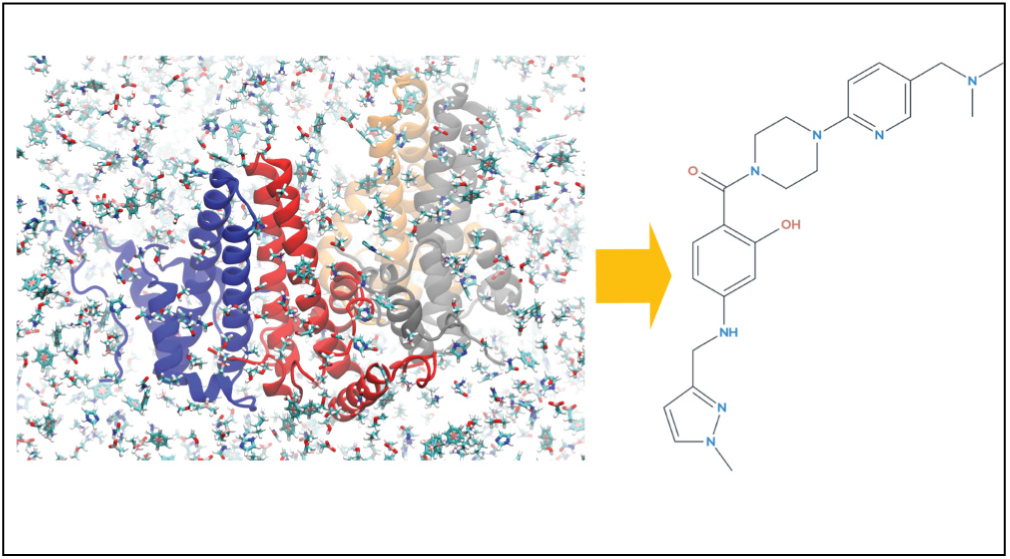

