## Supporting Information for "SILCS-Guided Feature Encoding Expands Ligand Recognition at an HBV Core Protein Interface"

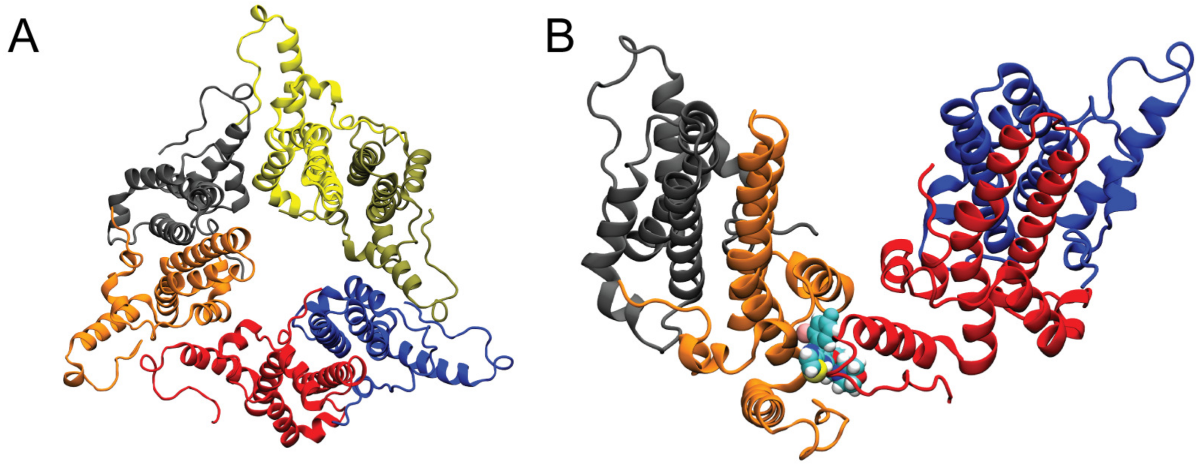


Figure S1: Structural context of the Cp149 hexamer and canonical capsid assembly modulator (CAM) binding site. **(A)** Ligand-free Cp149 hexamer shown in top view, corresponding to a trimer-of-dimers assembly. **(B)** Representative capsid assembly modulator (CAM) GLS4 bound at the canonical interdimer interface. GLS4 is shown in van der Waals representation.

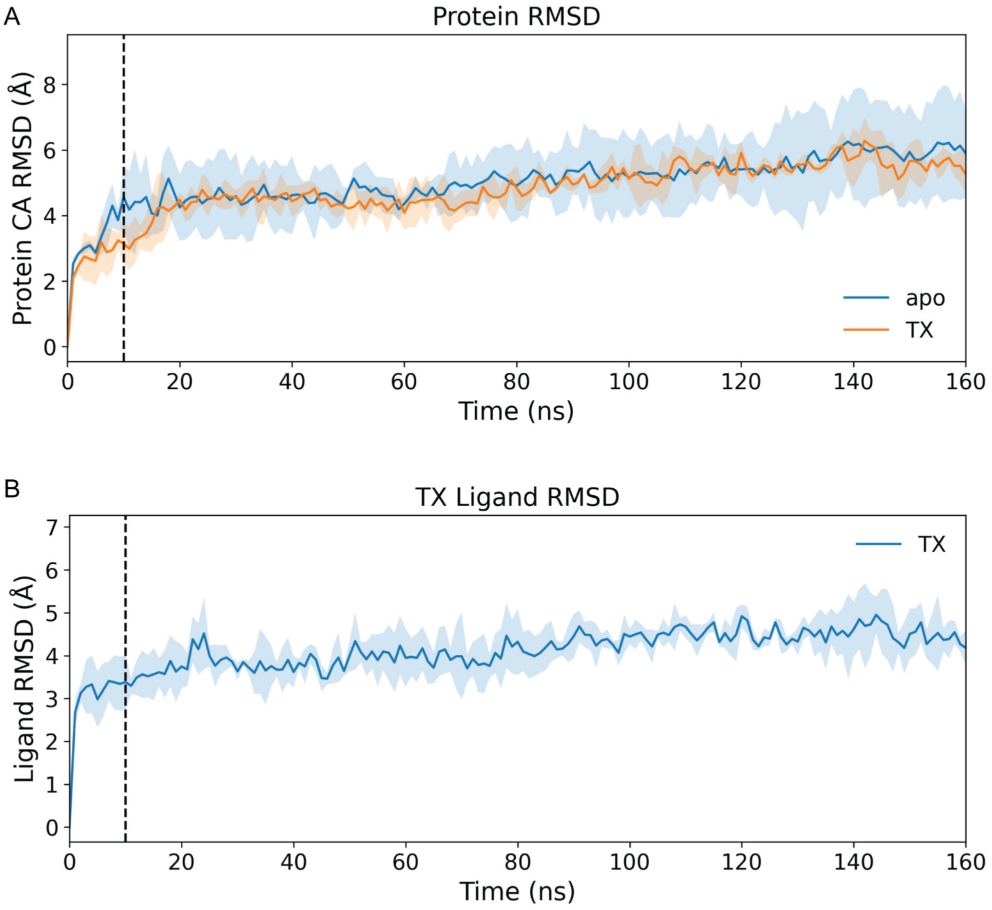


Figure S2: RMSD analysis of simulation stability across three independent 160-ns replicas. (A) Backbone $C_\alpha$ RMSD of the protein for the apo and TX100-bound systems, calculated relative to the respective initial frame. (B) RMSD of TX100 ligands in the bound simulations. In both panels, the solid line represents the mean across three replicas and the shaded region indicates the standard deviation. The dashed vertical line at 10 ns marks the equilibration period excluded from subsequent analyses.

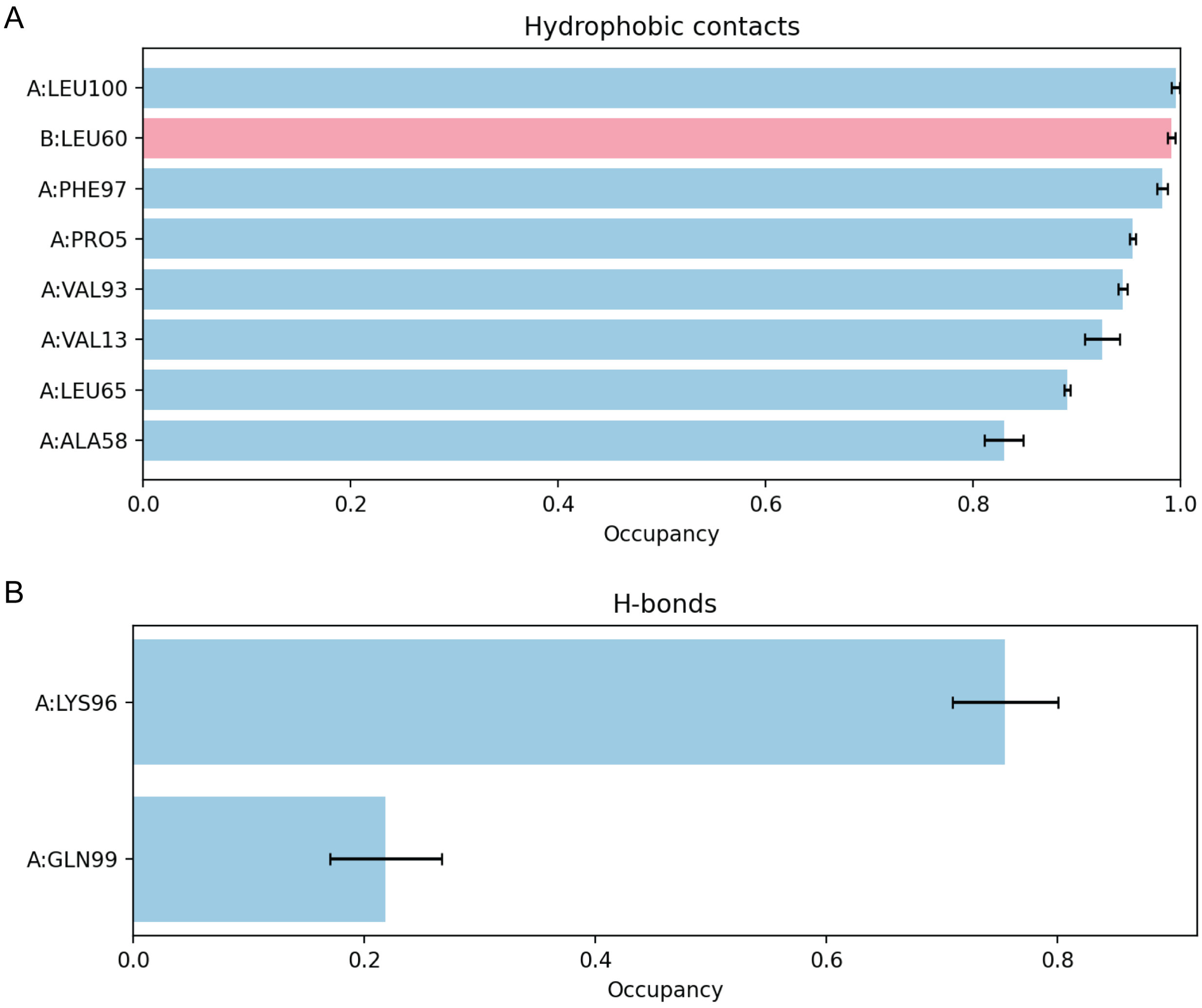


Figure S3: Quantitative analysis of TX100–protein interactions within the intradimer pocket. **(A)** Hydrophobic contact occupancies for residues interacting with TX100, calculated as the fraction of analyzed frames (excluding the first 10 ns) in which a contact is present. Only residues with average occupancy greater than 80% are shown. **(B)** Hydrogen-bond occupancies between TX100 and protein residues, calculated using the same trajectory window. Only residues with average occupancy greater than 10% are shown. Bars represent the mean occupancy across replicas, with error bars indicating the standard deviation. Bars are colored by chain, with chain A in light blue and chain B in light pink.

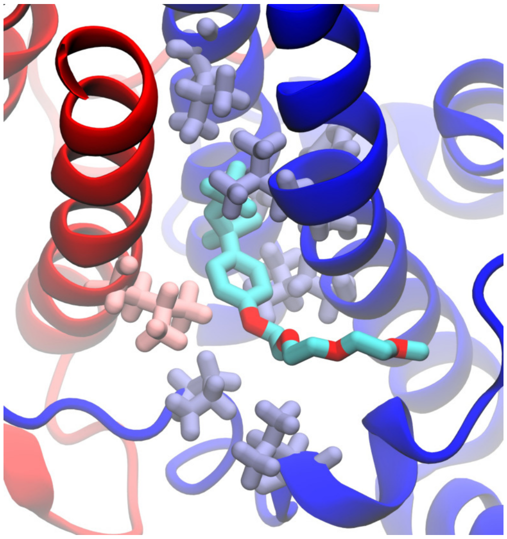

Figure S4: Structural view of high-occupancy hydrophobic contacts between TX100 and the Cp intradimer pocket. Chain A is shown in blue and chain B in red. TX100 and residues exhibiting persistent hydrophobic contacts with the ligand (> 80% occupancy) are shown in licorice representation. Contacting residues from chain A (Pro5, Val13, Ala58, Leu65, Val93, Phe97, and Leu100) are highlighted in ice blue, and the contacting residue from chain B (Leu60) is highlighted in pink. Hydrogen atoms are omitted for both ligand and residues.

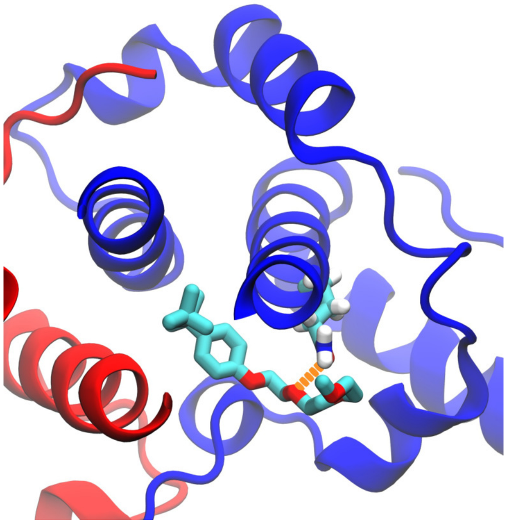

Figure S5: Representative hydrogen-bond interaction between Triton X-100 (TX100) and Gln99 at the Cp intradimer pocket. TX100 and Gln99 are shown in licorice representation, with hydrogen atoms on TX100 omitted. This interaction corresponds to the more transient polar contact observed near the pocket entrance.

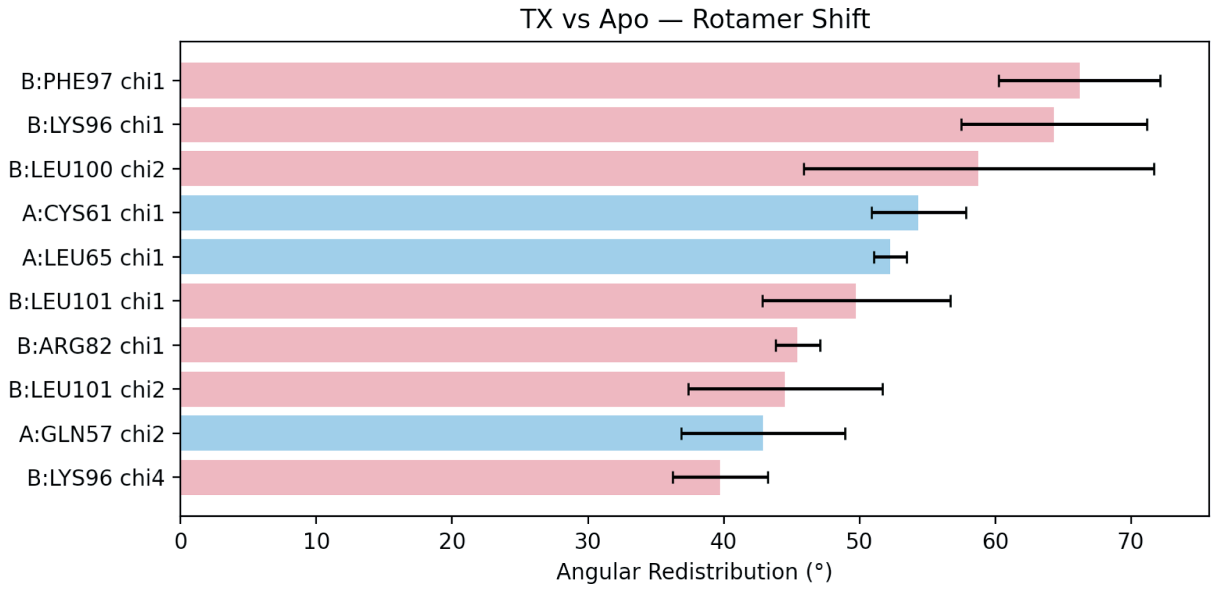


Figure S6: Rotamer redistribution of pocket-lining residues in the TX100-bound Cp149 hexamer relative to the apo system. Angular redistribution was quantified by comparing side-chain dihedral ($\chi$) angle distributions between the apo and TX100-bound simulations using a circular Wasserstein distance metric. Values represent the mean $\pm$ standard deviation across three independent replicas. Residues are shown only if the lower bound of the distribution (mean $-$ SD) exceeds 30°, indicating consistent and significant rotamer shifts. Residues are grouped by chain assignment, with chain A shown in blue and chain B shown in pink.

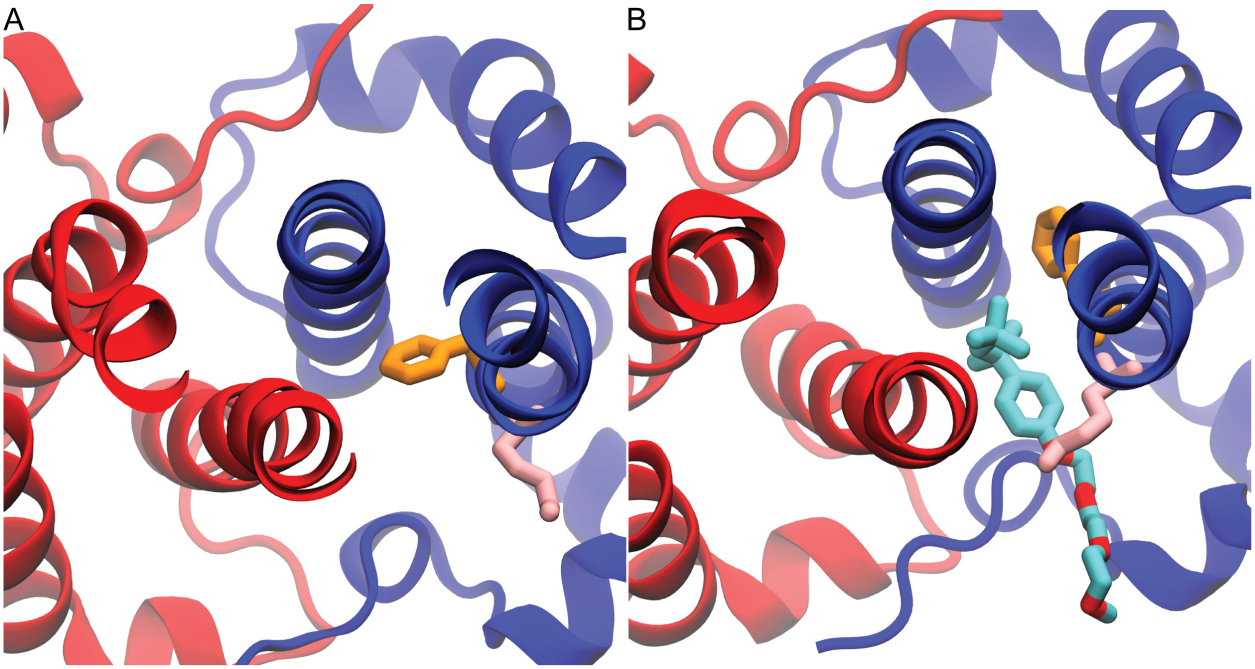


Figure S7: Representative side-chain conformations illustrating local structural heterogeneity of the Cp intradimer pocket in the apo and TX100-bound simulations. **(A)** Apo state, showing Lys96 and Phe97 from chain B in representative side-chain orientations. **(B)** TX100-bound state, showing ligand-associated rearrangement of Lys96 and Phe97, consistent with local pocket adaptability upon ligand binding. TX100 and residues Lys96 and Phe97 are shown in licorice representation, with Lys96 colored pink and Phe97 orange. Hydrogen atoms are omitted for both TX100 and the residues.

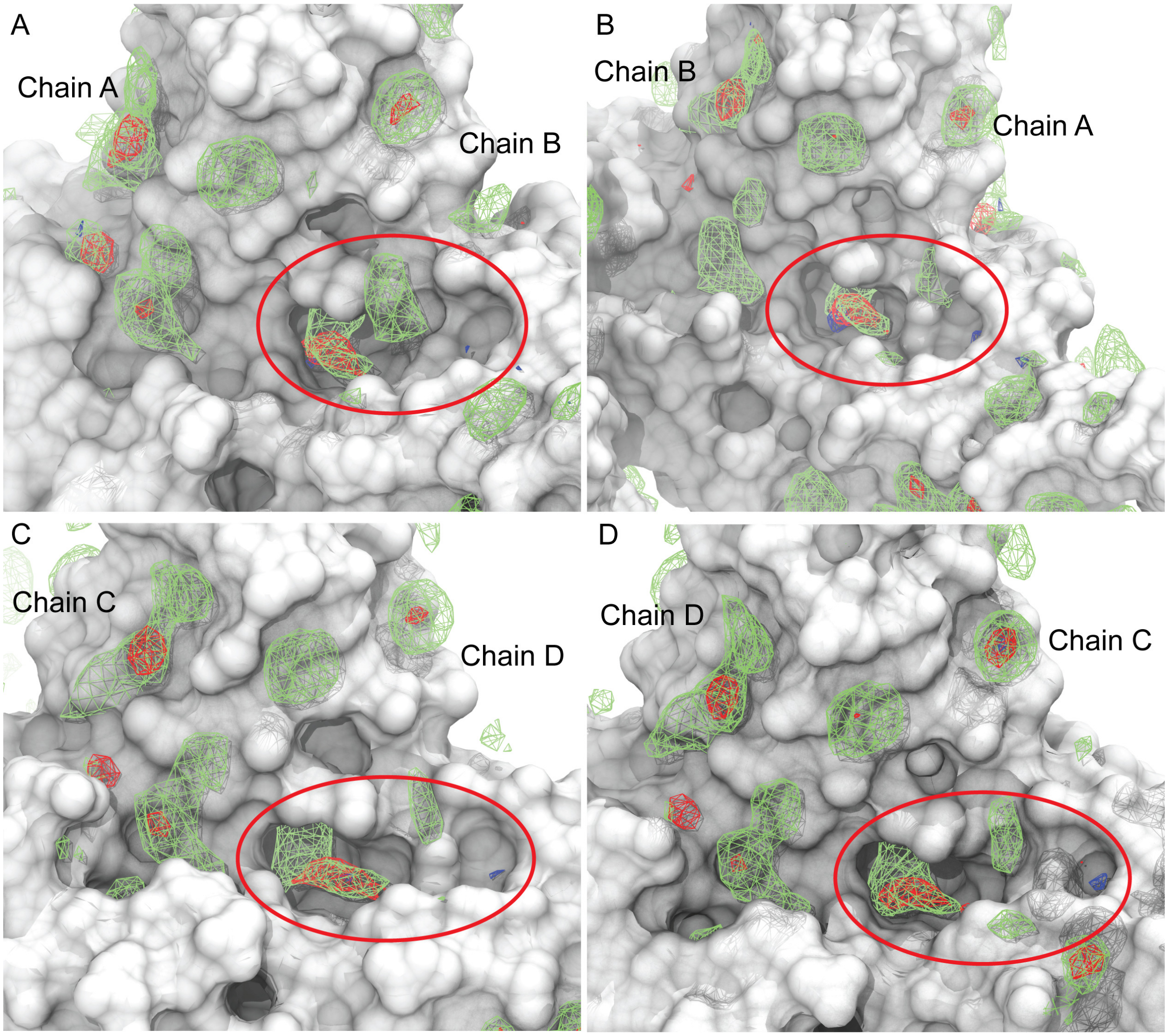


Figure S8: Comparison of SILCS FragMaps across the four intradimer pockets of the Cp149 tetramer. **(A,B)** Pockets formed between chains A and B on the exterior-facing (A) and interior-facing (B) sides. **(C,D)** Pockets formed between chains C and D on the exterior-facing (C) and interior-facing (D) sides. Apolar FragMaps are shown as green isosurfaces, hydrogen-bond acceptor features in red, and hydrogen-bond donor features in blue. The interior-facing A–B pocket shown in panel (B) was selected for subsequent pharmacophore construction and virtual screening.

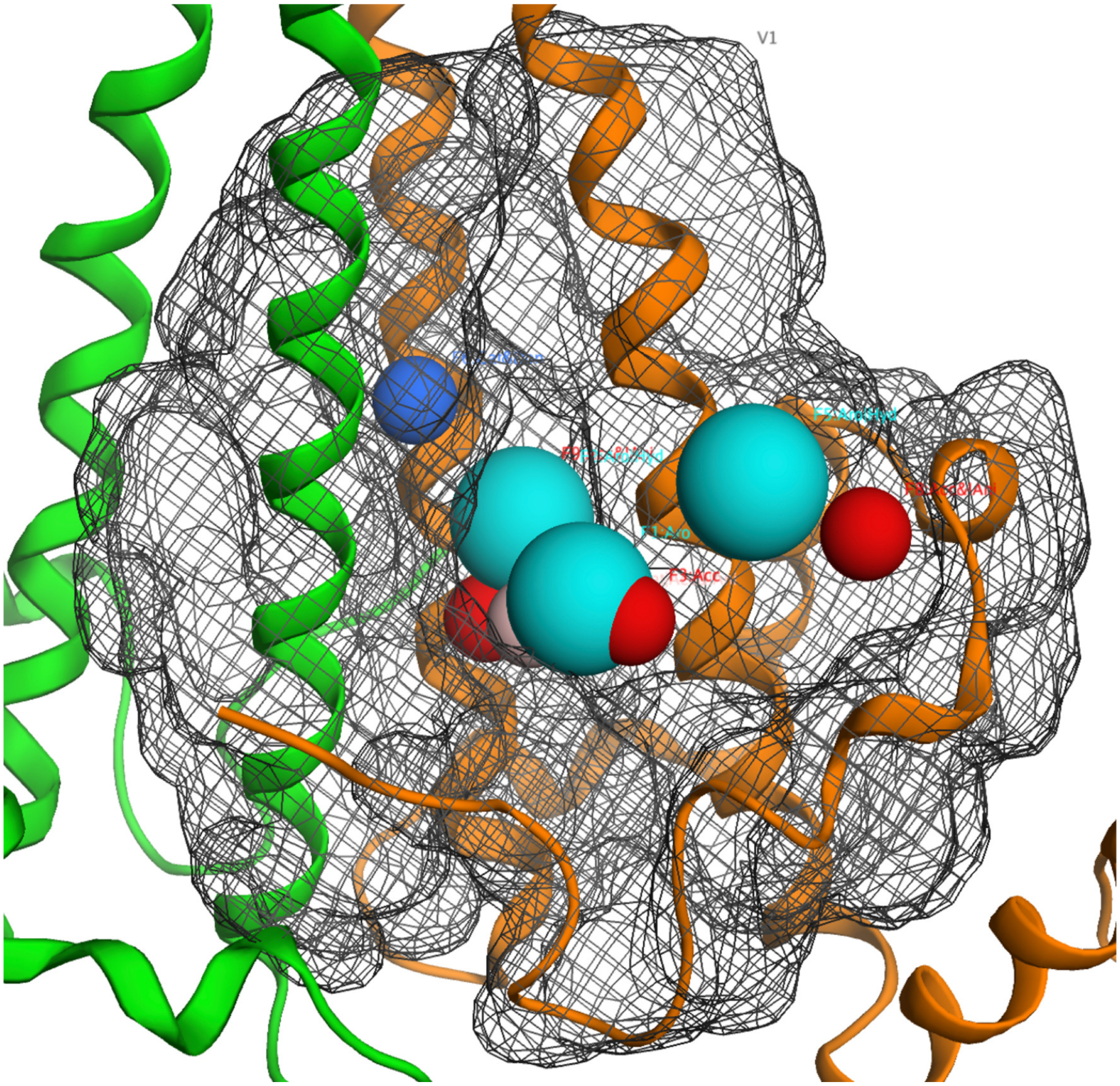


Figure S9: Initial pharmacophore feature identification from SILCS FragMap clustering in the selected intradimer pocket. FragMap voxels within the interior-facing A–B pocket were clustered based on spatial proximity and grid free energy favorability, yielding eight candidate interaction features. Chain A is shown in green and chain B in orange. Apolar features are shown as cyan spheres, hydrogen-bond acceptor features in red, and hydrogen-bond donor features in blue. Pink spheres indicate regions that can serve as either hydrogen-bond donors or acceptors, arising from the close spatial proximity of these two feature types. The grey grid surface represents the excluded volume, summarizing the space occupied by the protein during the simulation.

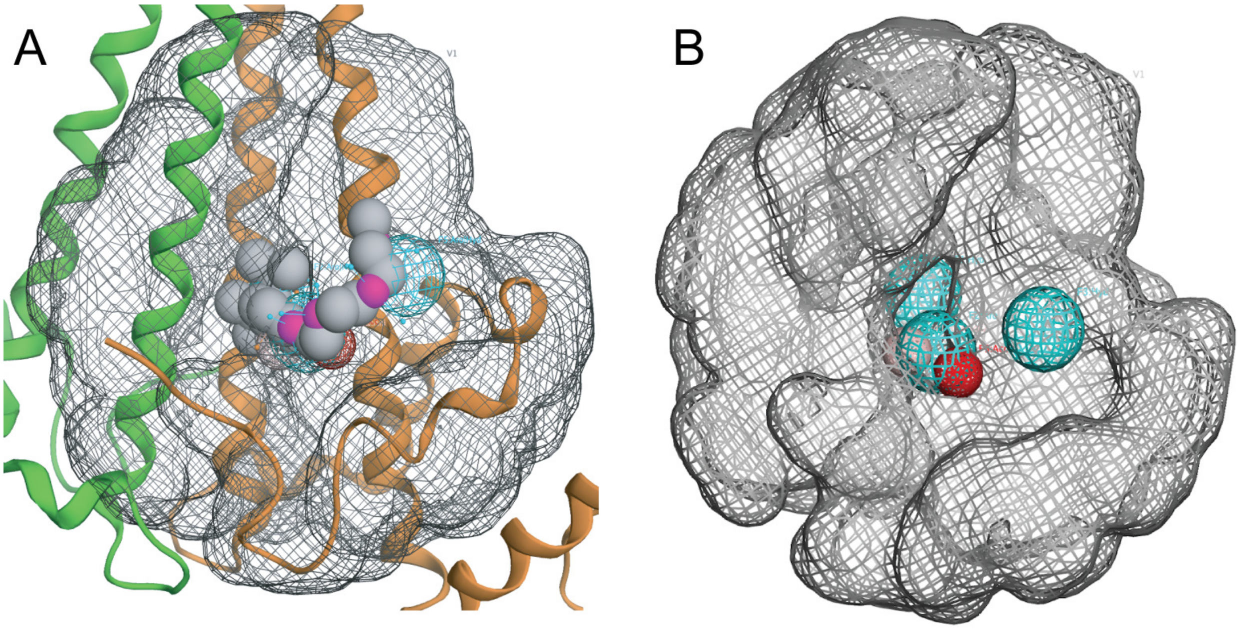


Figure S10: Final five-feature pharmacophore model and excluded-volume constraint used for virtual screening. **(A)** The final pharmacophore model corresponding to that shown in Fig. 2C, displayed within the Cp intradimer pocket with chain A in green and chain B in orange. The excluded-volume constraint is shown as a grey wireframe surface, and Features 1–5 are shown as colored spheres. **(B)** Isolated view of the final pharmacophore model and excluded-volume constraint, highlighting the spatial arrangement of the five selected interaction features within the sterically accessible region used for virtual screening.

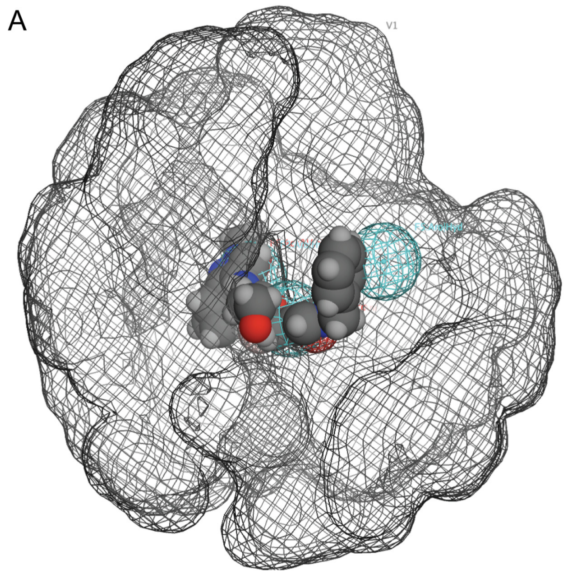


Figure S11: Representative examples of pharmacophore-based virtual screening results. Compounds are shown aligned to the pharmacophore model, illustrating how candidate molecules satisfy the required feature and geometric constraints used in the search. **(A)** Example for compound A4. **(B)** Example for compound B3.

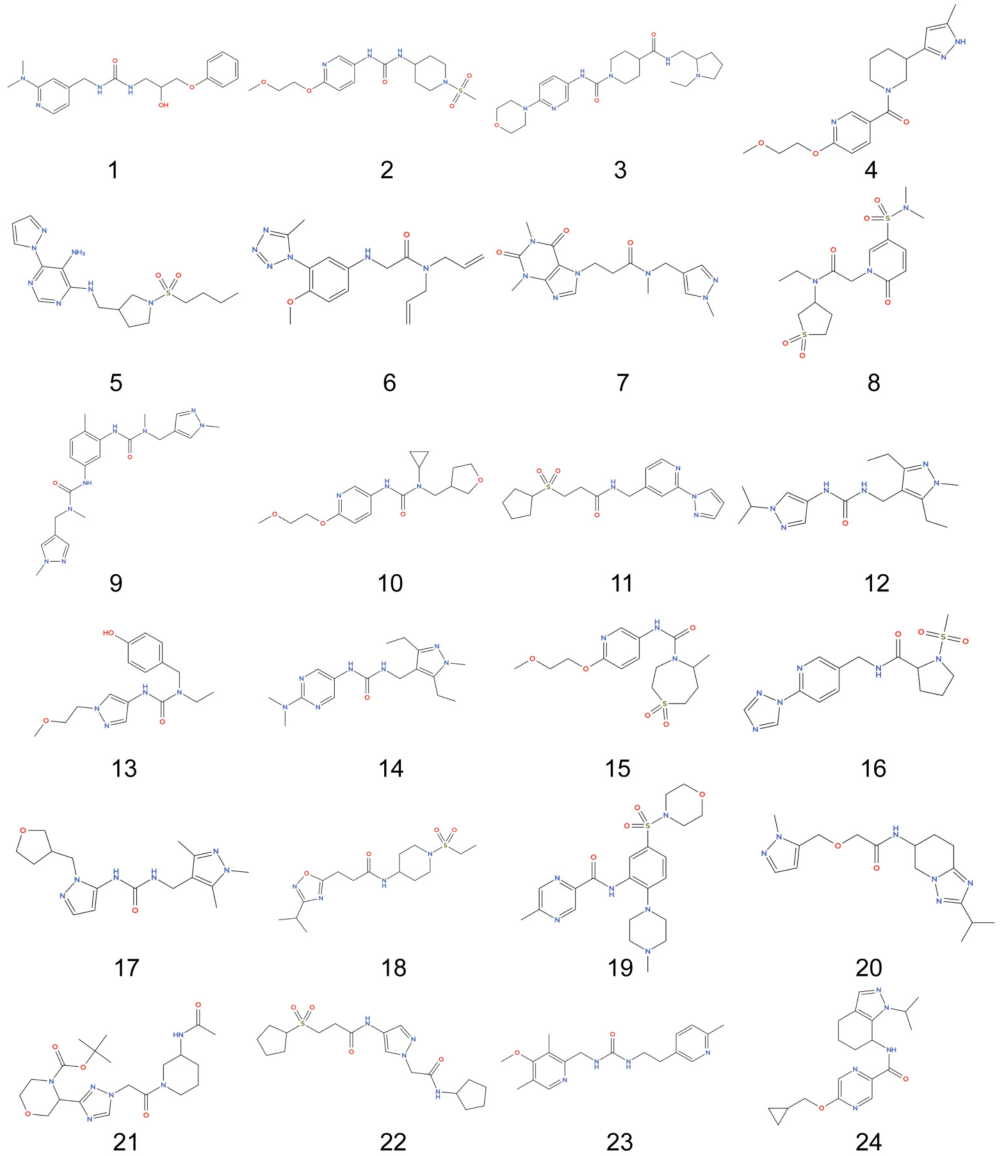


Figure S12: Chemical structures of candidate compounds (1–24) among the 120 molecules retained after molecular docking using a docking-score cutoff of −8 kcal/mol.

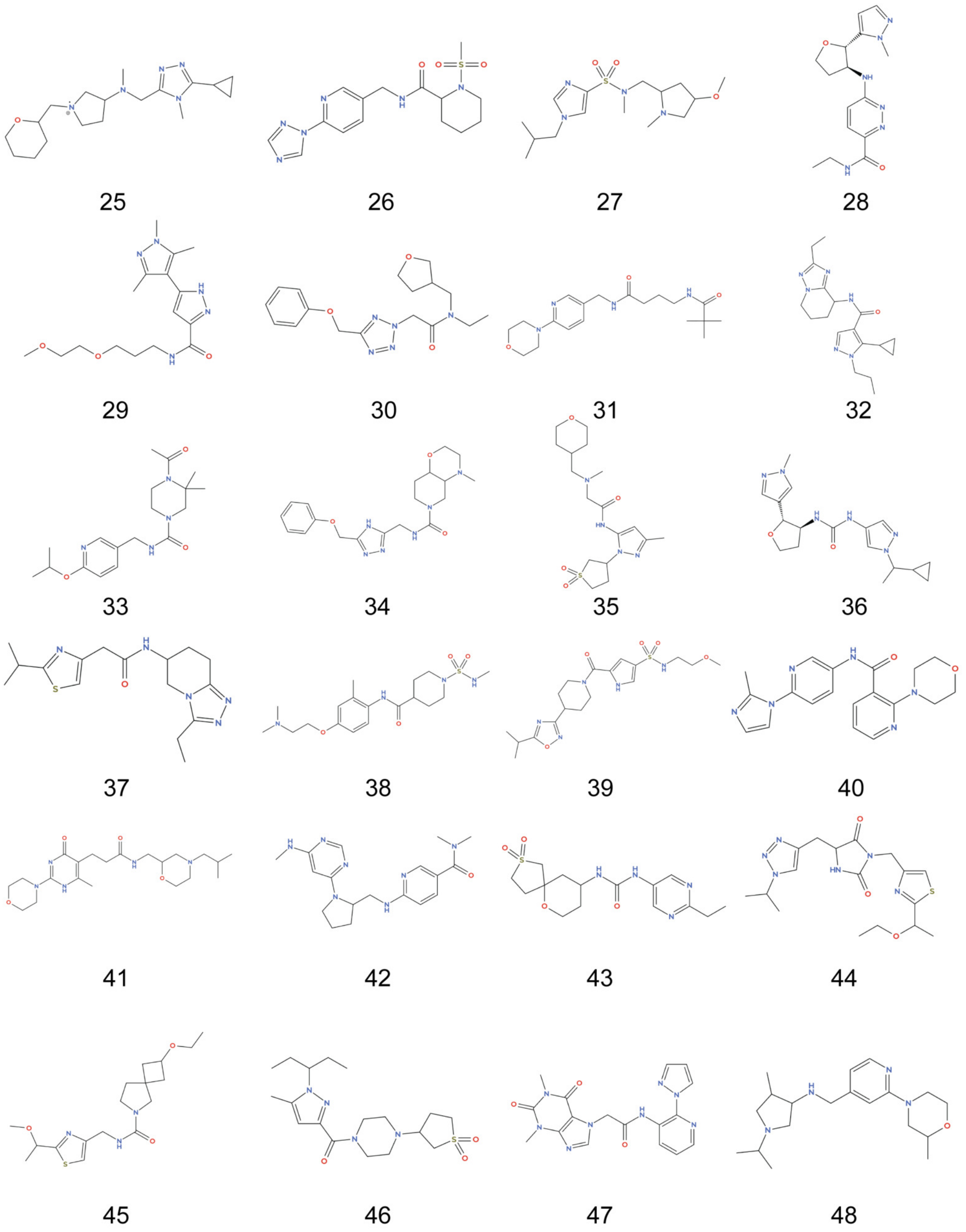


Figure S13: Chemical structures of candidate compounds (25–48) among the 120 molecules retained after molecular docking using a docking-score cutoff of −8 kcal/mol.

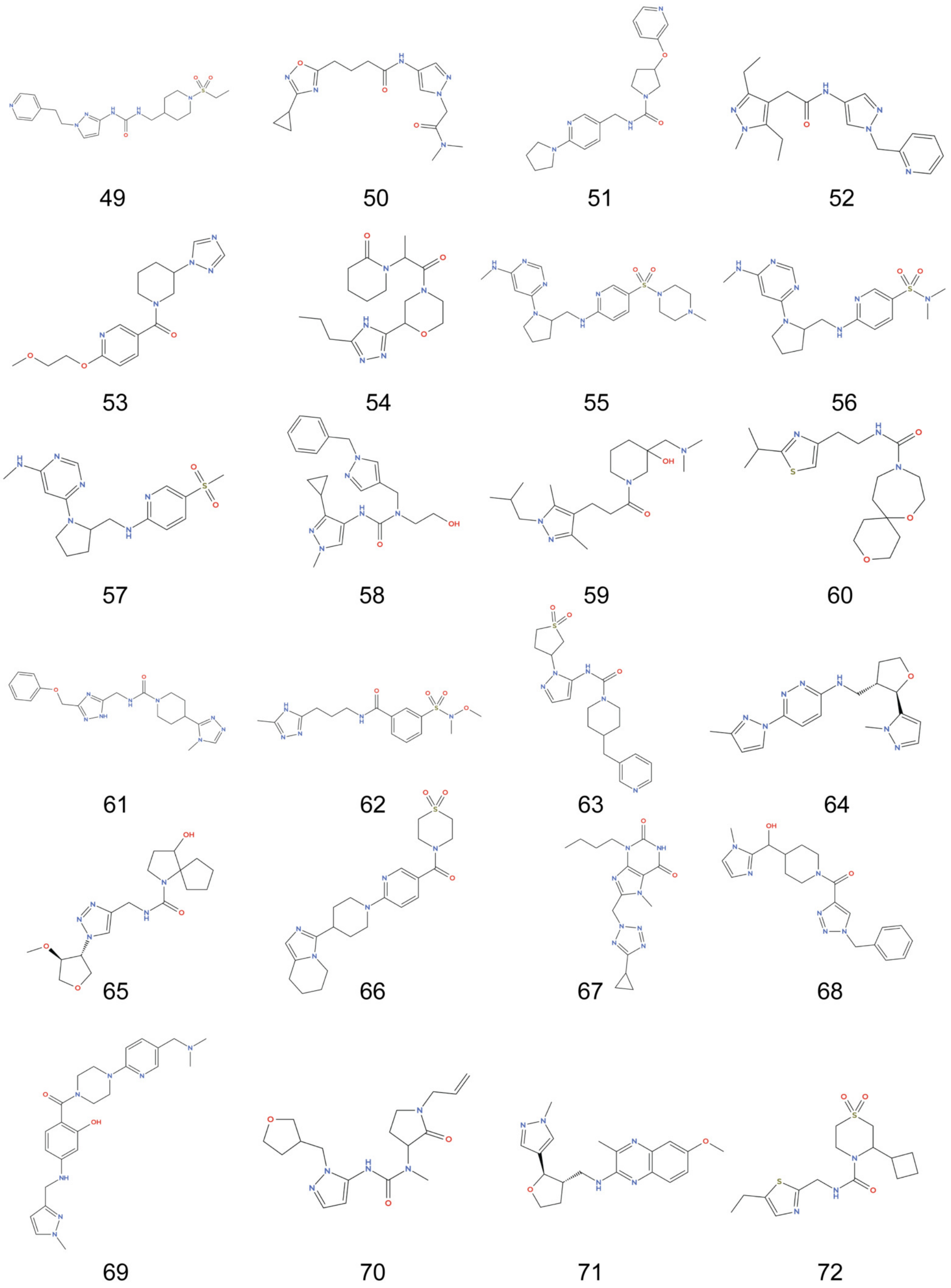


Figure S14: Chemical structures of candidate compounds (49–72) among the 120 molecules retained after molecular docking using a docking-score cutoff of −8 kcal/mol.

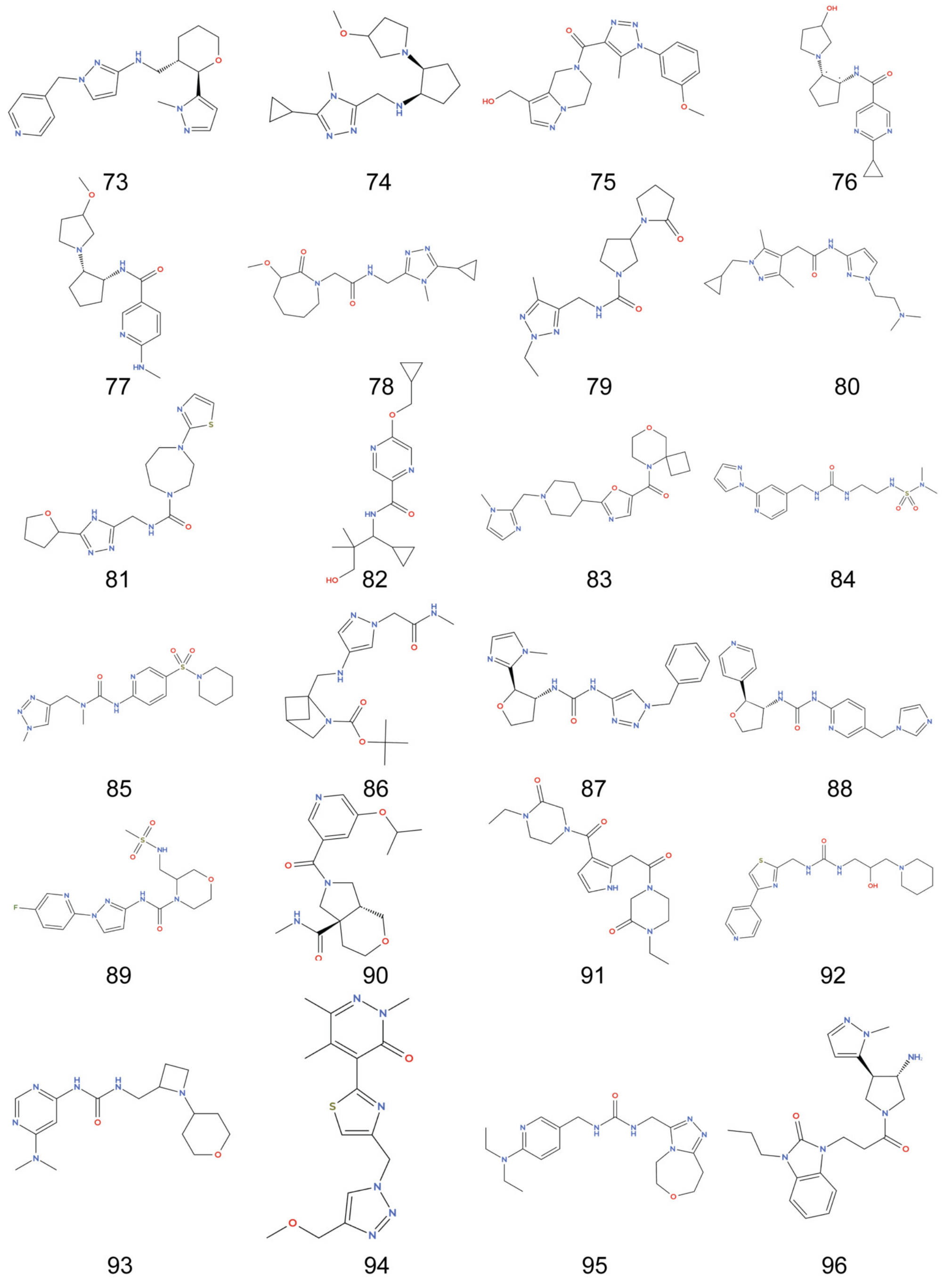


Figure S15: Chemical structures of candidate compounds (73–96) among the 120 molecules retained after molecular docking using a docking-score cutoff of −8 kcal/mol.

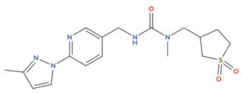
97

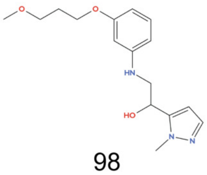
98

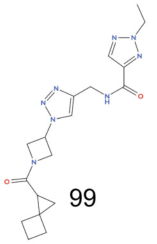
99

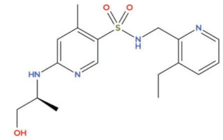
100

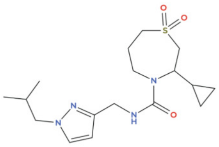
101

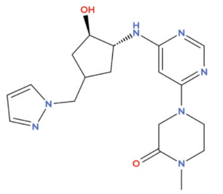
102

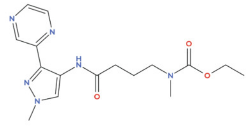
103

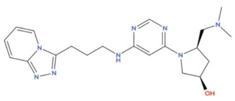
104

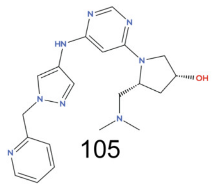
105

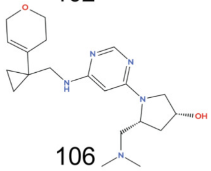
106

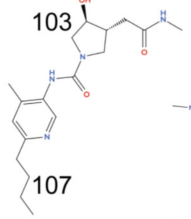
107

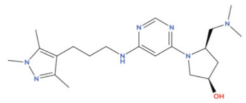
108

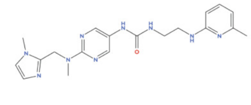
109

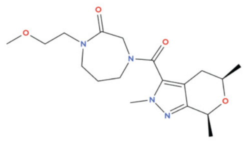
110

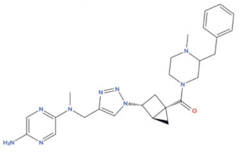
111

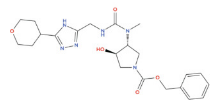
112

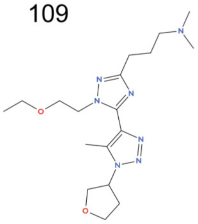
113

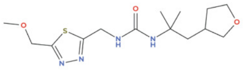
114

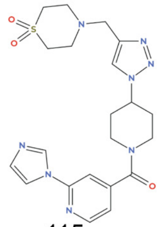
115

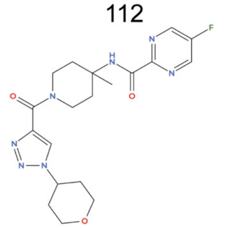
116

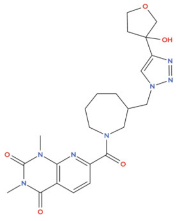
117

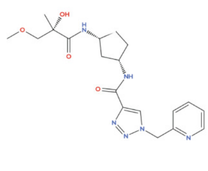
118

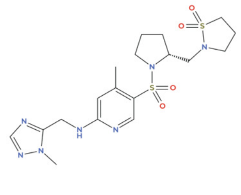
119

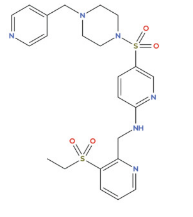
120

Figure S16: Chemical structures of candidate compounds (97–120) among the 120 molecules retained after molecular docking using a docking-score cutoff of −8 kcal/mol.

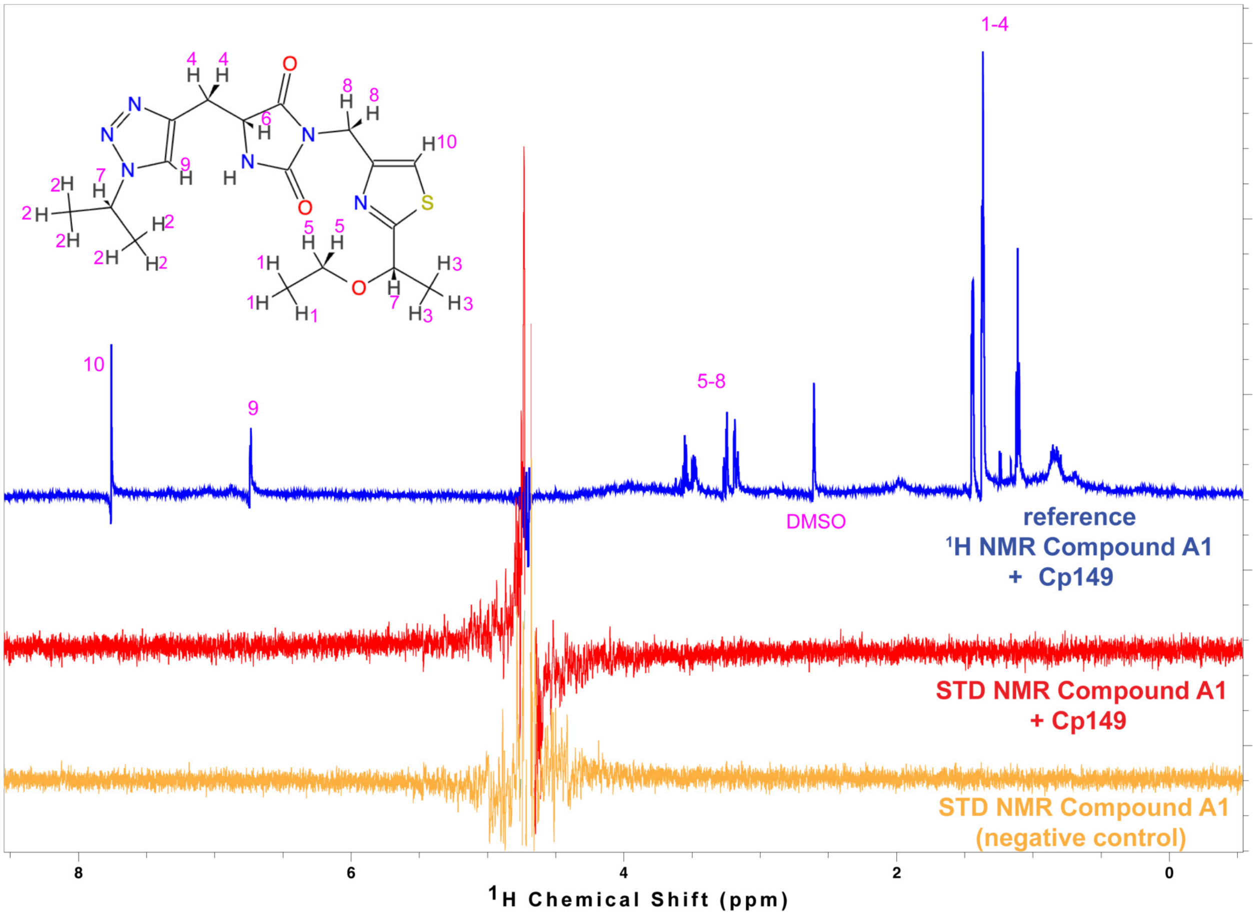


Figure S17: STD NMR characterization of ligand binding to Cp149 for compound A1.

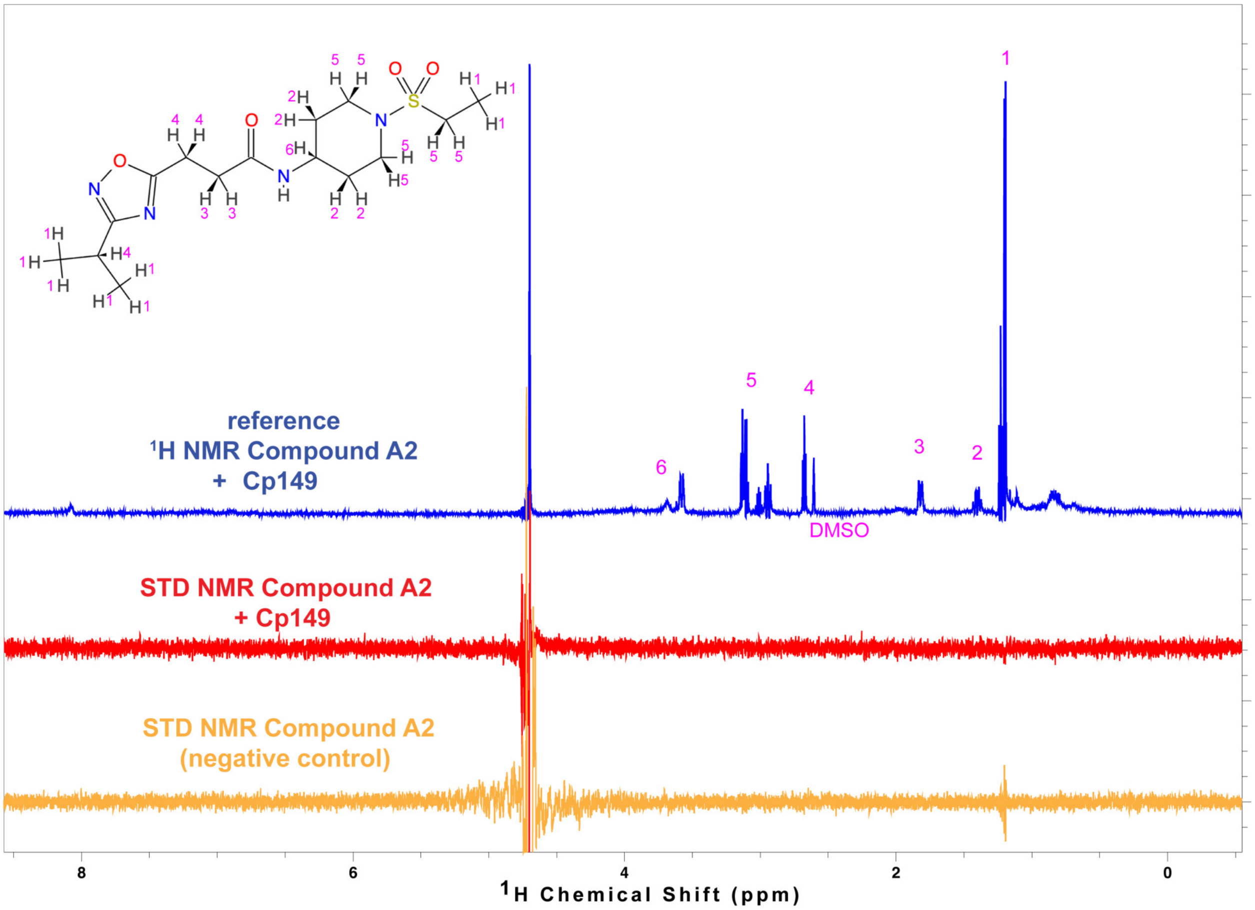


Figure S18: STD NMR characterization of ligand binding to Cp149 for compound A2.

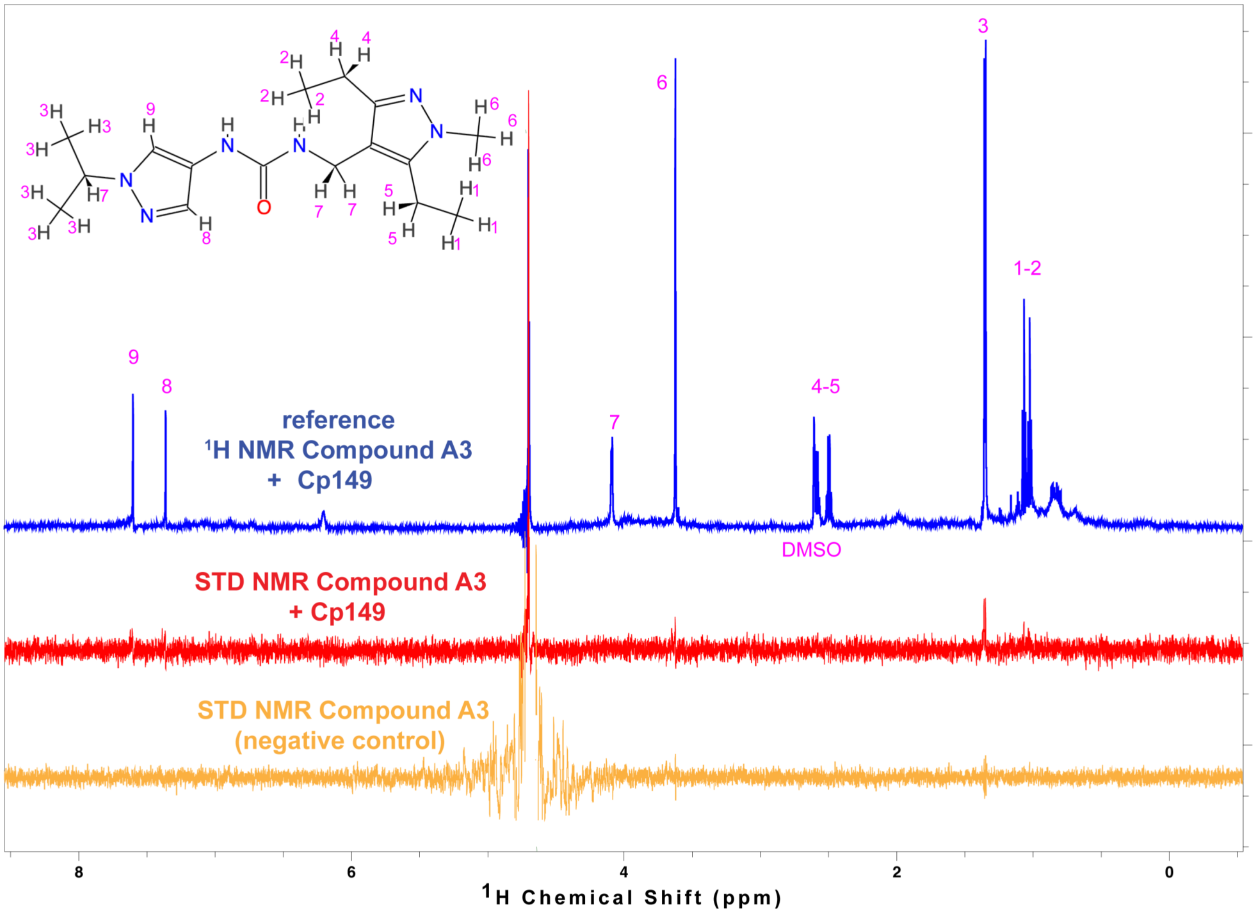


Figure S19: STD NMR characterization of ligand binding to Cp149 for compound A3.

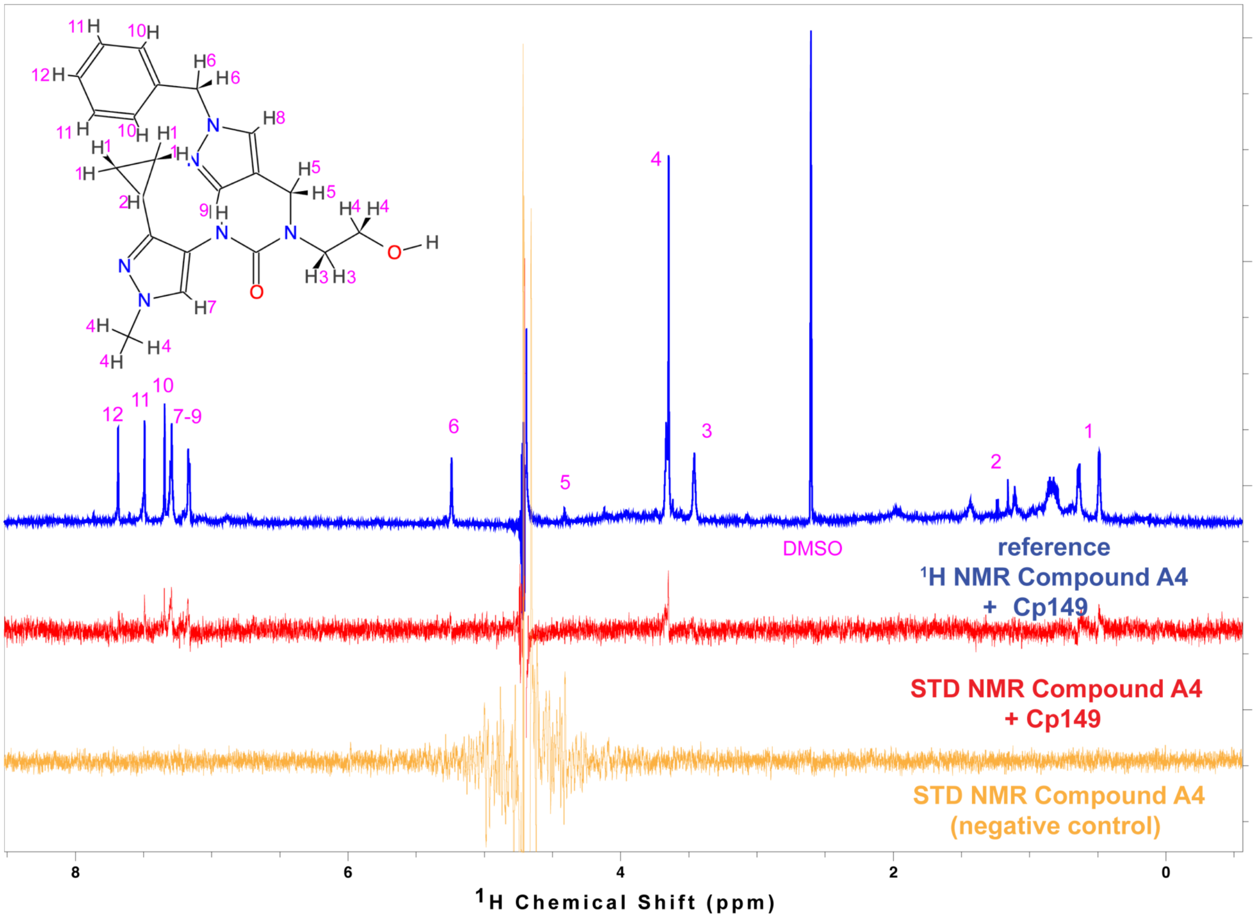


Figure S20: STD NMR characterization of ligand binding to Cp149 for compound A4.

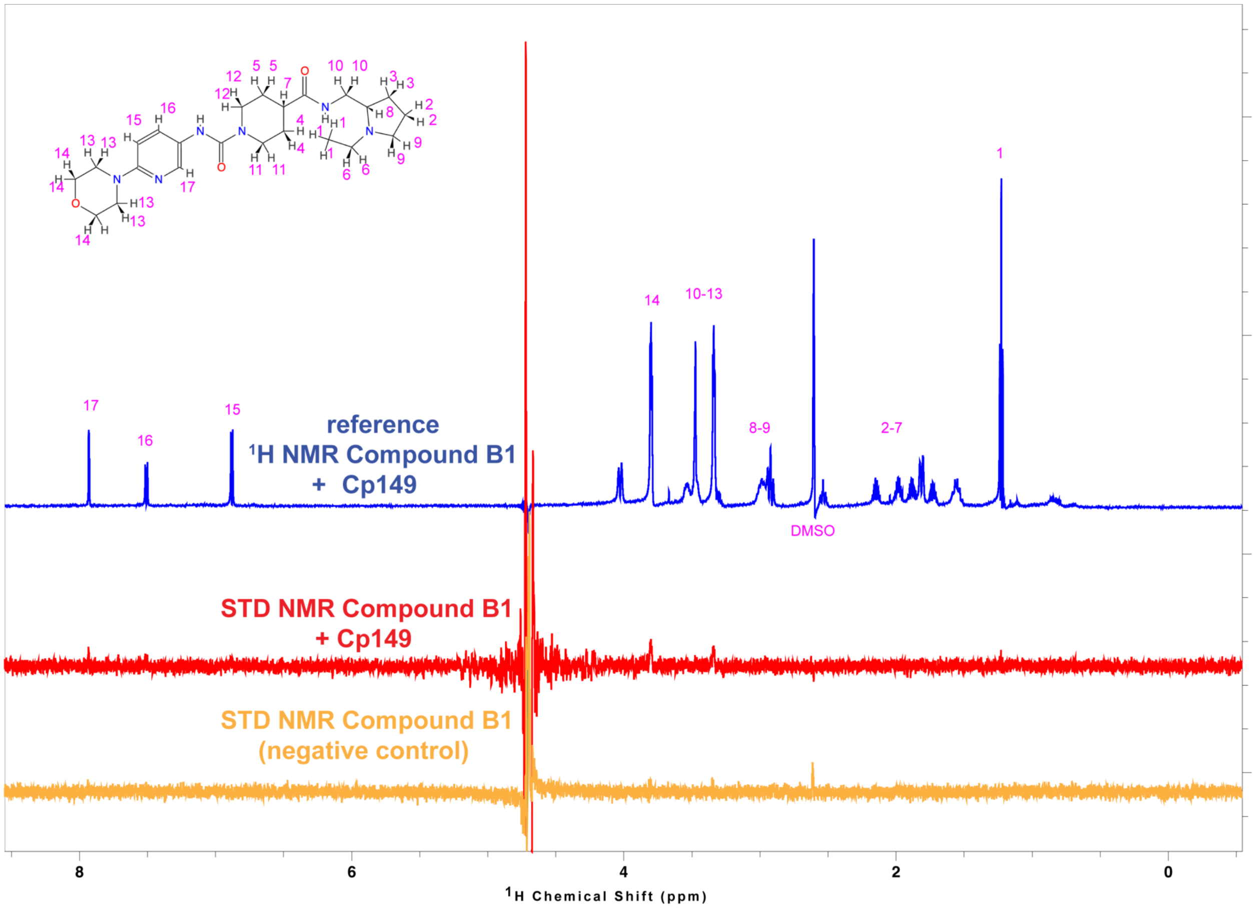


Figure S21: STD NMR characterization of ligand binding to Cp149 for compound B1.

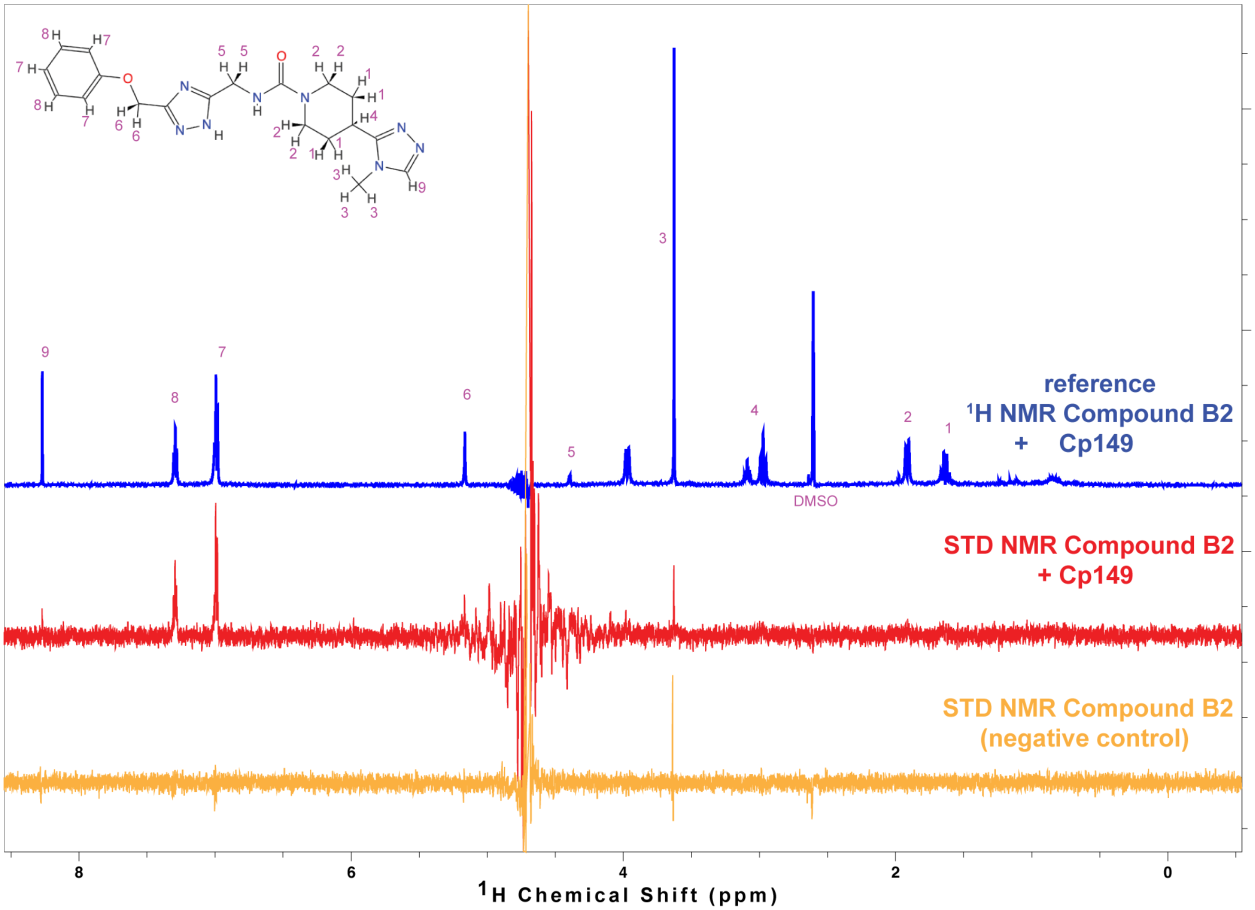


Figure S22: STD NMR characterization of ligand binding to Cp149 for compound B2.

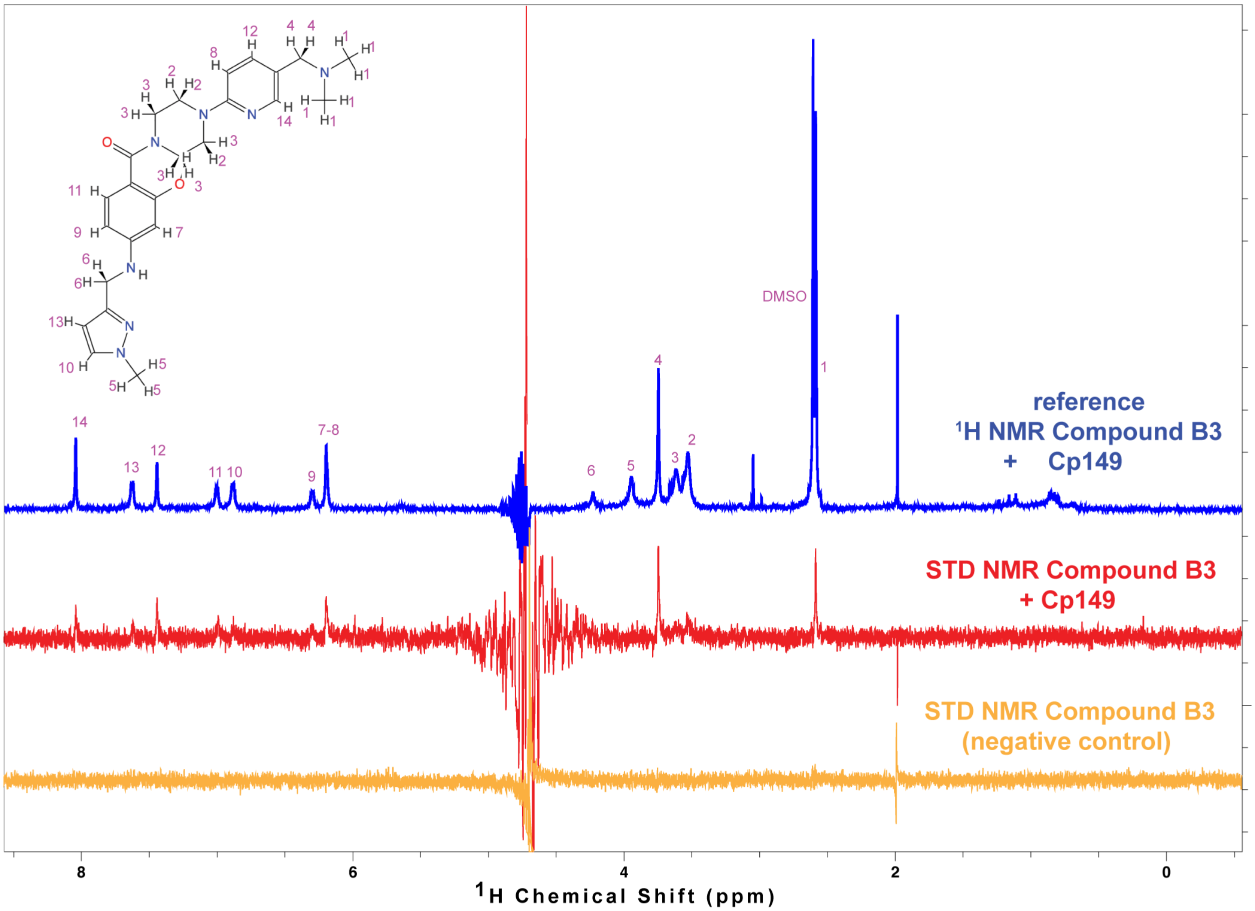


Figure S23: STD NMR characterization of ligand binding to Cp149 for compound B3.

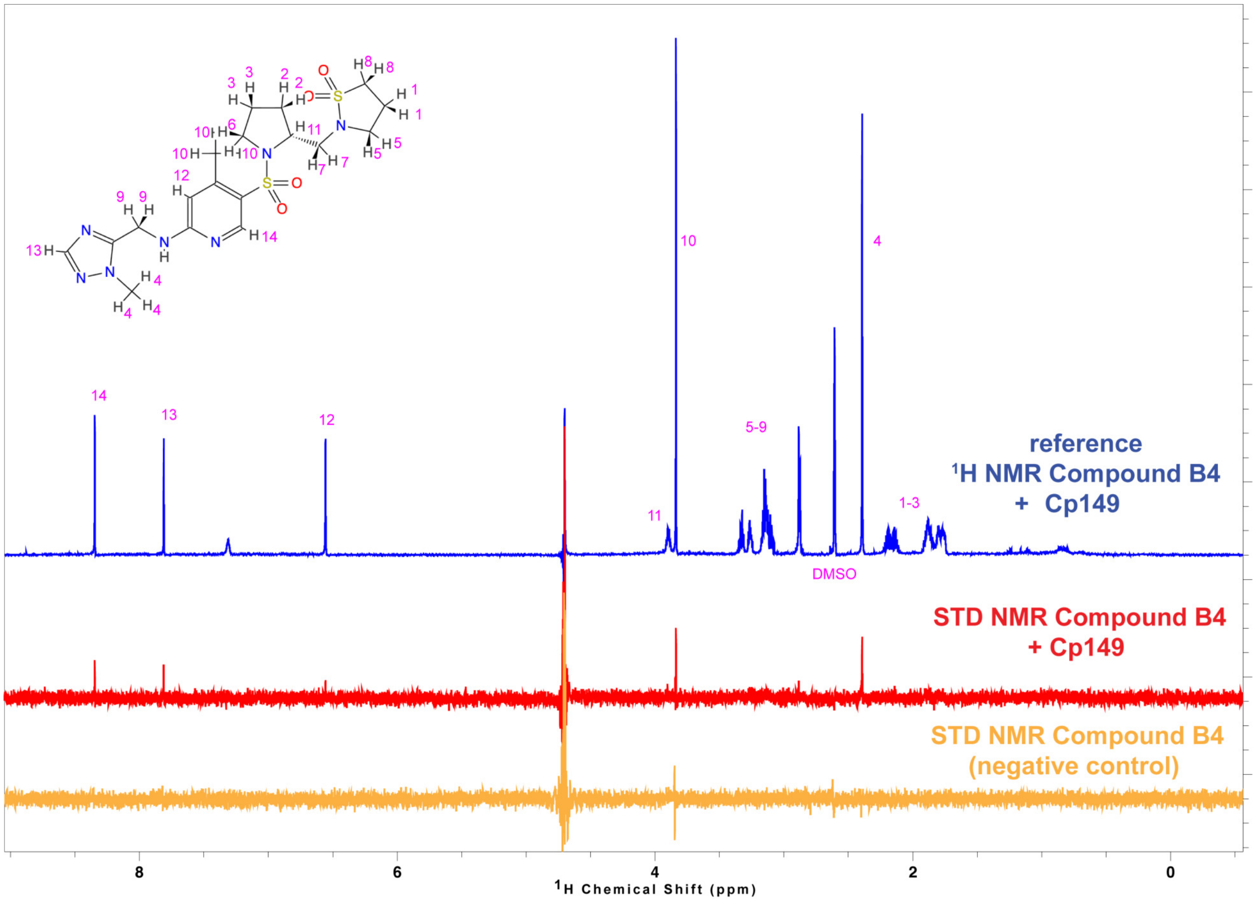


Figure S24: STD NMR characterization of ligand binding to Cp149 for compound B4.

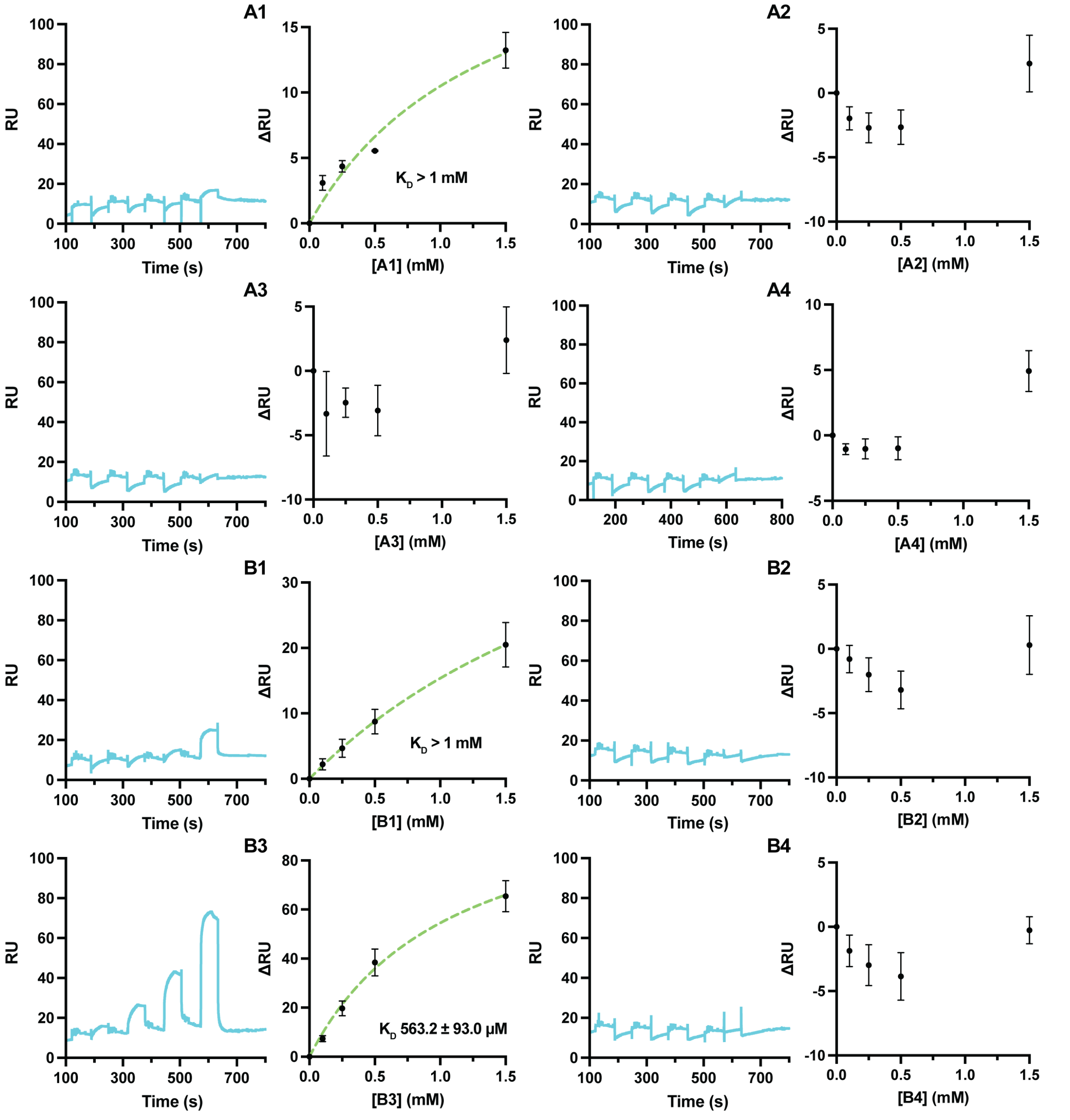


Figure S25: Surface plasmon resonance analysis of HBcAg-biotin binding to small-molecule ligands A1–B4. SPR measurements were performed using immobilized HBcAg-biotin with increasing concentrations of each compound (0, 0.1, 0.25, 0.5, and 1.5 mM). For each compound, SPR sensorgrams showing response units (RU) as a function of time are shown on the left, and steady-state changes in response units (ΔRU) as a function of ligand concentration are shown on the right. The $K_D$ determined from single-site binding model fitting for compound B3 is indicated. Data represent the mean ± standard deviation from three independent replicates.

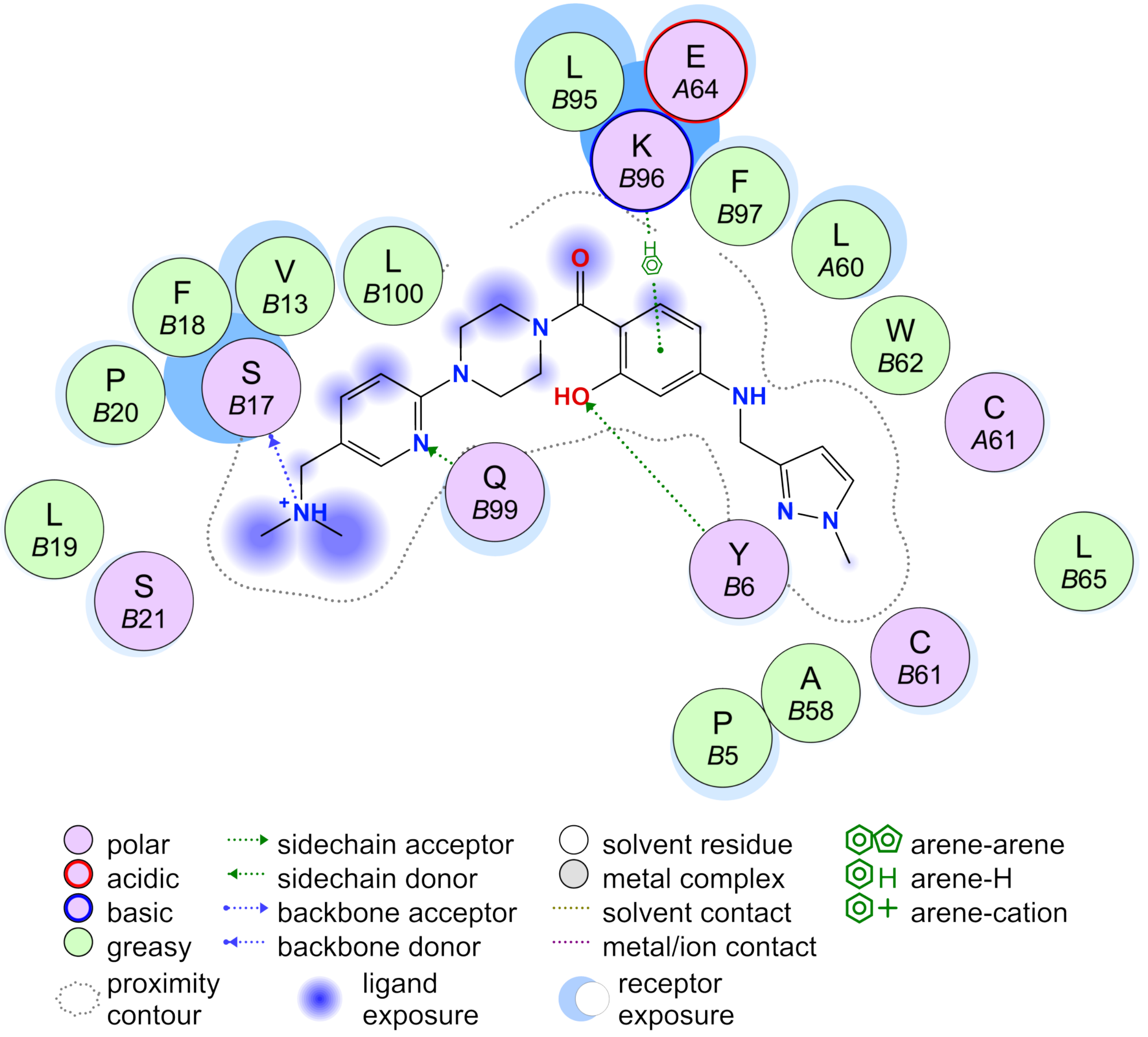


Figure S26: Legend for the two-dimensional ligand–protein interaction diagram shown in Fig. 4A. Symbols denote residue classes, hydrogen-bond interactions, hydrophobic contacts, solvent exposure, and ligand atom environments used in the interaction diagram.

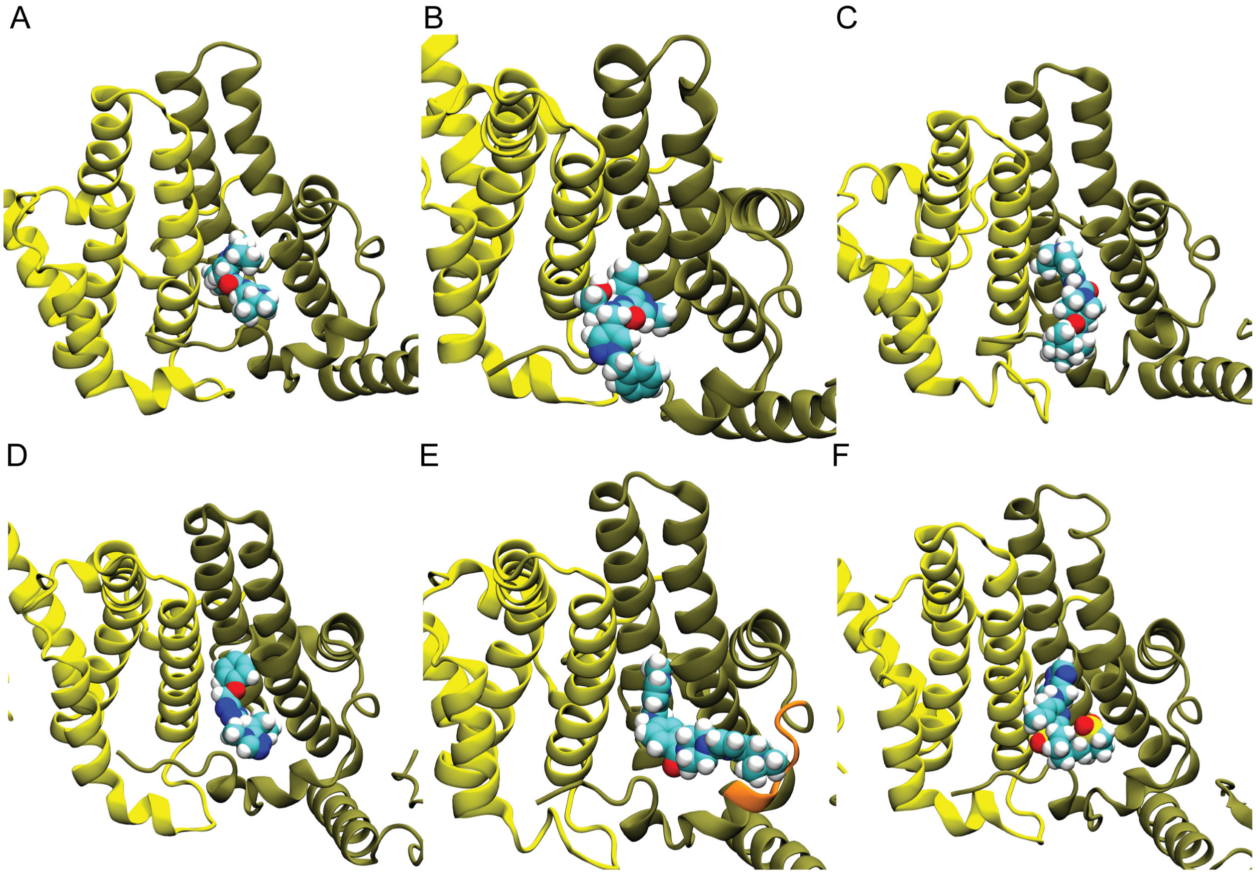


Figure S27: Representative binding modes from MD simulations for the six ligands exhibiting binding behavior. **(A)** A3, **(B)** A4, **(C)** B1, **(D)** B2, **(E)** B3, and **(F)** B4. In each panel, chain C is shown in yellow and chain D in tan, with ligands displayed in van der Waals representation. In panel **(E)**, the loop region of chain D (residues 18–22) is highlighted in orange, illustrating its interactions with ligand B3.

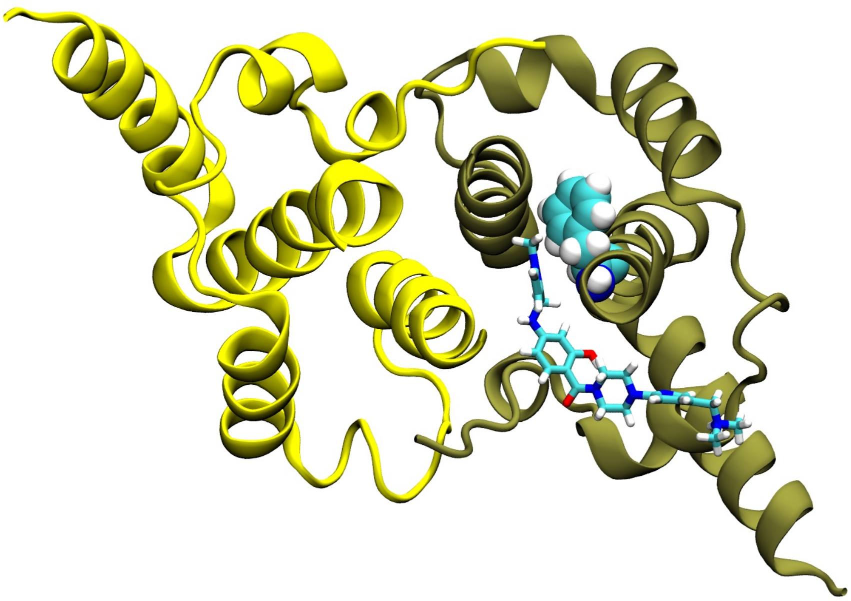

Figure S28: Representative Phe97 side-chain conformation in the B3-bound intradimer pocket. Chain C is shown in yellow and chain D in tan, with B3 displayed in licorice representation. Phe97 of chain D is shown in van der Waals representation.

Table S1: Filtering criteria applied during pharmacophore-based virtual screening

| **Property** | **Desired value** |
|---|---|
| Docking score | $< -8$ kcal/mol |
| Total atom count | 20–70 |
| Molecular weight | $< 500$ Da |
| Hydrogen bond donors | $\leq 5$ |
| Hydrogen bond acceptors | $\leq 10$ |
| H-bond donors + acceptors | $\leq 12$ |
| Rotatable bonds | $\leq 10$ |
| Topological polar surface area | $\leq 140$ Å$^2$ |
| Calculated LogP | $\leq 5$ |
| Molar refractivity | 40–130 |

Table S2: Docking scores and selected computed physicochemical properties of compounds chosen for experimental evaluation.

| **Compound** | **Docking score (kcal/mol)** | **TPSA (Å$^2$)** | **Molar refractivity** | **Rotatable bonds** |
|---|---|---|---|---|
| A1 | -8.77 | 102.24 | 99.14 | 8 |
| A2 | -8.60 | 105.40 | 88.91 | 8 |
| A3 | -8.47 | 76.77 | 90.43 | 8 |
| A4 | -8.42 | 88.21 | 109.47 | 10 |
| B1 | -8.71 | 91.24 | 123.86 | 9 |
| B2 | -8.18 | 113.85 | 103.94 | 8 |
| B3 | -8.48 | 90.96 | 128.39 | 8 |
| B4 | -8.20 | 130.39 | 114.19 | 7 |

Table S3: Summary of MD behavior across independent simulations for candidate ligands

| **Compound** | **Trial 1** | **Trial 2** | **Trial 3** |
|---|---|---|---|
| A1 | Unstable | Unstable | Unstable |
| A2 | Unstable | Unstable | Unstable |
| A3 | Localized | Localized | Localized |
| A4 | Unstable | Unstable | Unstable |
| B1 | Localized | Localized | Localized |
| B2 | Localized | Localized | Localized |
| B3 | Extended | Extended | Extended |
| B4 | Localized | Localized | Localized |

Localized indicates maintenance of a confined pocket-bound pose; extended indicates maintenance of a pocket-bound pose extending toward the spike-proximal loop; unstable indicates loss of the proposed pocket-bound pose during simulations. A1 and A2 dissociated during the initial equilibration window and are therefore not shown in the representative binding-pose figure.

Table S4: Summary of experimental and MD support for candidate ligands

| **Compound** | **STD NMR** | **SPR** | **MD behavior** |
|---|---|---|---|
| A1 | No obvious signal | Weak response, $K_D > 1$ mM | Unstable |
| A2 | No obvious signal | No appreciable response | Unstable |
| A3 | Weak/ambiguous | No appreciable response | Localized |
| A4 | Detectable signal | No appreciable response | Unstable |
| B1 | Weak/ambiguous | Weak response, $K_D > 1$ mM | Localized |
| B2 | Detectable signal | No appreciable response | Localized |
| B3 | Detectable signal | $K_D = 563.2 \pm 93.0$ $\mu$M | Extended |
| B4 | Detectable signal | No appreciable response | Localized |

STD NMR classifications are based on comparison of ligand–Cp149 samples with ligand-only controls. SPR classifications are based on concentration-dependent binding to immobilized HBcAg-biotin. MD behavior classifications are defined as follows: localized indicates maintenance of a confined pocket-bound pose with limited engagement beyond the hydrophobic cavity; extended indicates maintenance of a pocket-bound pose extending toward the spike-proximal loop; unstable indicates loss of the proposed pocket-bound pose.
